# A photo-switchable yeast isocitrate dehydrogenase to control metabolic flux through the citric acid cycle

**DOI:** 10.1101/2021.05.25.445643

**Authors:** Haoqi Chen, Lianne Mulder, Hein J. Wijma, Ronja Wabeke, Jose Pedro Vila Cha Losa, Mattia Rovetta, Tijn Caspar de Leeuw, Andreas Millias-Argeitis, Matthias Heinemann

## Abstract

For various research questions in metabolism, it is highly desirable to have means available, with which the flux through specific pathways can be perturbed dynamically, in a reversible manner, and at a timescale that is consistent with the fast turnover rates of metabolism. Optogenetics, in principle, offers such possibility. Here, we developed an initial version of a photo-switchable isocitrate dehydrogenase (IDH) aimed at controlling the metabolic flux through the citric acid cycle in budding yeast. By inserting a protein-based light switch (LOV2) into computationally identified active/regulatory-coupled sites of IDH and by using *in vivo* screening in *Saccharomyces cerevisiae*, we obtained a number of IDH enzymes whose activity can be switched by light. Subsequent *in-vivo* characterization and optimization resulted in an initial version of photo-switchable (PS) IDH. While further improvements of the enzyme are necessary, our study demonstrates the efficacy of the overall approach from computational design, via *in vivo* screening and characterization. It also represents one of the first few examples, where optogenetics were used to control the activity of a metabolic enzyme.

## Introduction

Metabolism is a complex network, with its enzymes being regulated at many different levels. Intricate system behavior can arise from such regulation, for instance, biostability during startup of glycolysis^1^, or metabolic oscillations^2–4^. Furthermore, metabolism is tightly intertwined with other cellular functions. For instances, it was recently found that it also interacts with the eukaryotic cell cycle, possibly in a system of coupled oscillators^5^. Metabolism was also found to regulate transcription through chromatin acetylation with the highly regulated metabolite acetyl-CoA^6,7^.

To study the complexity of metabolic systems, and to disentangle the connections to other cellular functions, experimental tools are required that allow us to dynamically perturb metabolism in a targeted manner. While recent microfluidic technology enables dynamic changes in nutrient supply^8,9^, such perturbations are confined to the respective entry-points of the nutrients. In contrast, with enzyme inhibitors also reactions further away from the entry-points can be perturbed. However, inhibitors can suffer from slow uptake dynamics^10^ and potential off-target effects^11^. Alternatively, gene and protein levels could be perturbed with either chemical inducers^12^ or optogenetic means^13^, or targeted protein depletion^14^. While with these means, i.e., the induction of protein expression or protein depletion, in principle, every metabolic enzyme can be perturbed, these are rather slow perturbations compared to the time scale at which metabolism operates. Thus, in essence, we still lack experimental tools, with which we can perturb metabolism in a highly dynamic, reversible and enzyme-targeted manner, ideally in single cells.

One emerging possibility to achieve such perturbation is provided by optogenetics. Optogenetics is a technique that exploits light-sensitive protein domains, which alter their three-dimensional structure upon exposure to light of a specific wavelength. Optogenetic protein domains have been used to induce protein oligomerization^15^, translocation^16^, or transcription^13,17^. Analogously, it should be feasible to fuse optogenetic domains to catalytically active proteins, and to transduce light-induced changes in the optogenetic domain into structural changes of the protein to alter its activity. This is, in fact, something that nature has already accomplished, for instance, with the light-activatable adenylate cyclase in the soil bacterium Beggiatoa^18,19^, where blue light illumination reversibly activates cAMP production via a BLUF (blue light receptor using FAD) domain. Indeed, recent work has demonstrated that such functionality can also be engineered. For instance, to control the activity of certain protein kinases by light, an engineered light-sensitive domain, pdDronpa^20^, the small photo-switchable domain LOV2 (Light, Oxygen and Voltage protein domain 2)^21^, or the recent Light-Regulated domain engineered from Vivid photoreceptor^22^, were integrated into the kinase in a manner, such that light altered its activity. The same was recently also accomplished for the metabolic enzymes, e.g. the *Escherichia coli* dihydrofolate reductase^23^ or the mammalian pyruvate kinase^24^.

Here, given its importance for being at the cross-road between the oxidative part of the tricarboxylic acid (TCA) cycle, the glyoxylate shunt and the nitrogen assimilation metabolism, we developed a first version of a photo-switchable NAD^+^-dependent isocitrate dehydrogenase in *S. cerevisiae*. Exploiting the natural mechanism of its allosteric regulation^25^, and adopting a recent computational method^21^, we designed and constructed a library with IDH variants, performed *in-vivo* screening, characterized and optimized variants obtained from the screening. We hope that after further improvement, the engineered photo-switchable IDH variant will be used to address new research questions on the control of TCA cycle activity.

## Result

### Design of insertion of optogenetic domain

The NAD^+^-specific *Saccharomyces cerevisiae* IDH is a hetero-octamer; in specific a tetramer of heterodimers consisting of a regulatory (IDH1, 360 aa) and a catalytic subunit (IDH2, 369 aa)^25^. The regulatory IDH1 subunits form the core of the hetero-octamer, whereas the IDH2 subunits are positioned at the periphery of the enzyme (Fig. 1A). Both IDH1 and IDH2 domains have a substrate binding pocket that binds isocitrate, as well as a nucleotide binding pocket, which on IDH1 binds the allosteric regulator AMP but on IDH2 binds the cofactor NAD+^25^ (Fig. 1A). Upon coordinated binding of isocitrate and AMP on IDH1, structural transitions are propagated via the interacting surface between the IDH1/IDH2 hetero-dimer to increase the IDH2 affinity for isocitrate and therefore the catalytic rate of the enzyme^25,26^.

**Fig. 1:**
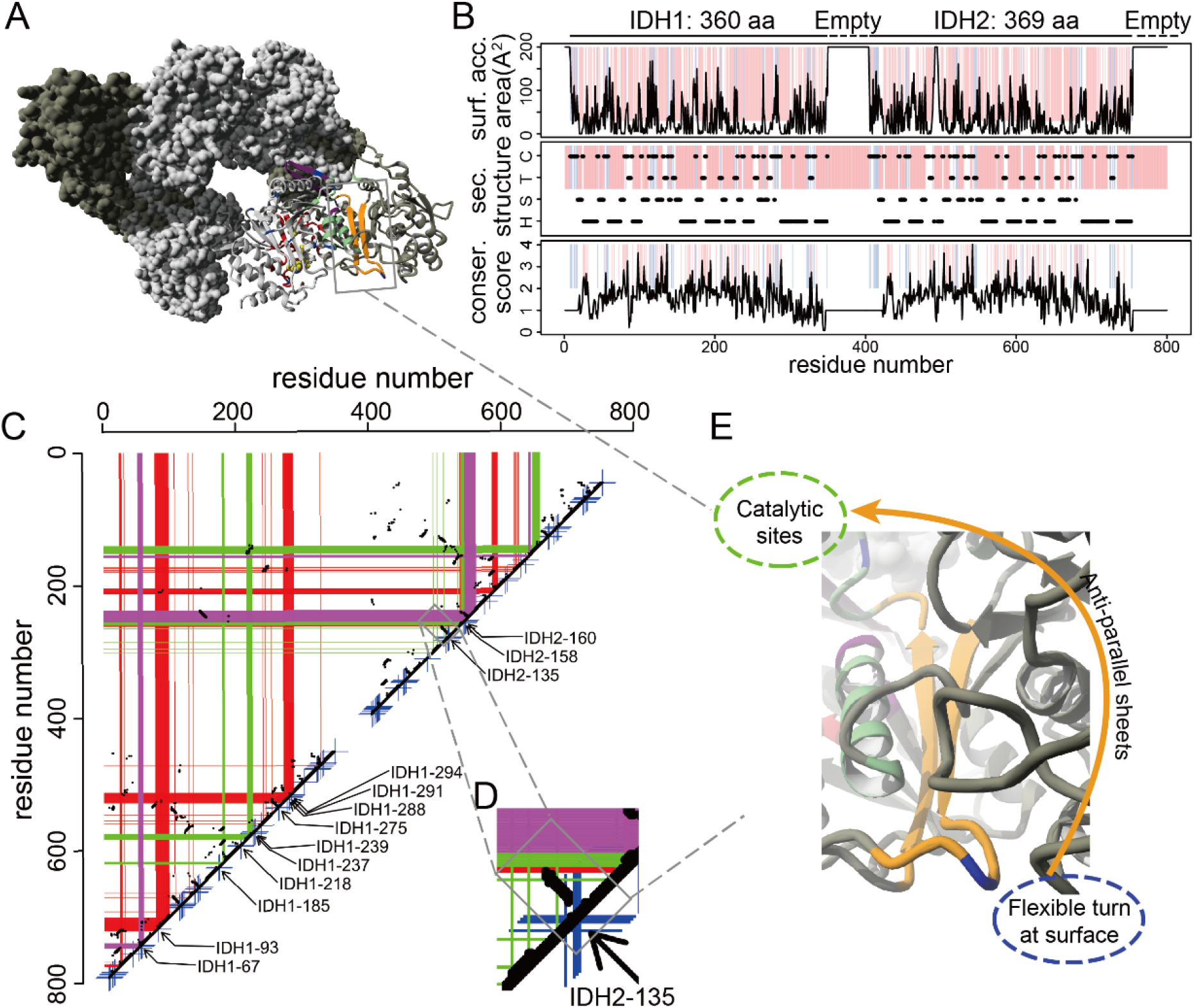
Computational identification of suitable LOV2 insertion site in IDH1 and IDH2. (A). Full structure of IDH octamer, bound with citrate (isocitrate analog), AMP and NAD+ (yellow beads). IDH1 domains are colored in light grey while IDH2 domains are colored in dark grey. Residues involved in catalysis are colored in green, residues in the allosteric binding site for AMP and citrate are colored in red, while residues in the allosterically altered regions are colored in purple. The structure is adapted from PDB structure 3blx. (B). Selection of positions suitable for LOV2 insertion based on sufficient surface accessible area (defined as solvent exposure) (upper panel), secondary structure (with turn/coil selected) (middle panel), and a Kullback-Leibner conservation score lower than 2 (lower panel). The x-axis is adjusted so that IDH1 falls on 0-359 aa and IDH2 falls on 400-769 aa, to aid visualization. In each panel, black curves or dashed lines plot the corresponding values on the y-axis; the residues that were filtered out by the current criteria were colored in red. After filtering with all three criteria, 106 residues remained and are colored in blue in all panels. Abbreviations for the middle panel: H, α-helical structure; B, β-sheet; T, turn; C, coil. (C). Contact map of IDH. The axes are the same as in (B), i.e., IDH1 falls on 0-359 and IDH2 on 400-769. Black dots correspond to Cα-Cα contacts within 7 Å that occur at least once within the hetero octamer. Short blue segments that cross at the diagonal indicate positions where insertions are allowed according to the criteria in (B). Green bands correspond to the catalytic sites, red bands correspond to the allosteric binding sites, and purple bands correspond to the allosterically altered regions25, as shown in the structure (A). (D). An example showing a suitable insertion site on the contact map. In this example, the IDH2-135 residue has passed all three criteria as indicated in (B) and is also connected to the active site 248-259 and allosteric binding site 138-139 (residue positions on 3blx), with a series of contacts less than 7 Å. The actual structure of the flexible loop containing IDH2-135 can be seen at (E). (E). The coil (blue, residues IDH2-135/136/137) is connected with the catalytic sites via a pair of anti-parallel sheets. The LOV2 domain will be inserted between IDH2-135 (residue indicated in blue) and IDH2-136.

Towards our aim to alter the IDH’s enzyme activity by light, we had to identify a suitable site in either of the two protein subunits, where we could insert the optogenetic domain LOV2, whose structural changes when induced by light would have an effect on the IDH enzyme activity. Inserting the 143-aa LOV2 domain into either of the two IDH subunits would result in four LOV2 domains to be presented on the holoenzyme. The insertion should meet two criteria: First, it should largely preserve the native conformation of IDH, allowing either the dark or the light-exposed state of the IDH-LOV2 fusion protein to be catalytically active. Second, structural changes in the helices of the inserted LOV2 domains, induced by light, should propagate to the IDH active or regulatory sites to in turn induce structural changes that lead to changes in catalytic activity of the IDH. To identify sites that satisfy these criteria, we modified and applied a computational pipeline that has recently been applied on several protein kinases^21^.

First, to identify sites where LOV2 insertion would lead to minimal perturbation of the native IDH conformation, for each residue on the IDH holoenzyme, we determined (i) the surface accessible area, (ii) the secondary structure using YASARA^27^, and (iii) the Kullback-Leibner conservation score characterizing the evolutionary conservation of a residue^28^ (Fig. 1B). On the basis of these features, we selected residues (i) with an accessible surface area of >30 squared angstrom (Å^2^), (ii) that fall within a turn/coil as the respective flexibility facilitates insertion of the optogenetic domain, and (iii) that have a conservation score of less than 2, as less conserved residues tend to be less important for enzyme functioning^29,30^. In total, we identified 106 residues that fulfill all three criteria, located in 31 turns and 12 coils.

Second, to further narrow down residues that would allow for propagation of light-induced structural changes in LOV2 into structural changes in IDH, we computed a residue ‘contact map’ (Fig. 1C), where we show which two residues within the holoenzyme are less than 7 Å away from each other (based on Cα-Cα distance in the protein structure), as done recently^21^. A series of contacts that extends perpendicularly from the diagonal (Fig. 1D) or are located away from the diagonal usually represent a tight loop comprising of anti-parallel helixes or sheets (Fig. 1E), which connects adjacent structures (in the peptide chain) or remotely located structures (in the 3D structure)^21^. If such tight loop connects a regulatory/catalytic region with a turn/coil that fulfill the above criteria, then it could allow propagation of light-induced structural changes (Fig. 1E), and the corresponding turn/coil would be further selected.

We then superimposed the regulatory/catalytic regions as well as the 31 turns and 12 coils selected above of IDH on this map (Fig. 1C), and searched for such tight loops as well as the corresponding turns/coils for LOV2 domain insertion. Some turns/coils that contain regulatory/catalytic residues were also selected (Fig. S1). In total, we selected 15 turns/coils (Table S1). By visually examining each site, we excluded two turns, which were positioned such that LOV2 insertion should have caused steric clashes with the rest of the protein. In case a turn/coil contains more than one residue, we selected the one that is most proximal to the N-terminus as insertion site, and inserted the LOV2 domain behind the selected residues (proximal to the C-terminus). This procedure resulted in total in 13 possible insertion sites, with three located in IDH2 and ten in IDH1 (Table S1). We used CRISPR/Cas9 (Ref. 31) to insert the LOV2 domain at the respective position in the *Saccharomyces cerevisiae* genome.

### *In vivo* screening for light-inducible variants

To identify candidates of the constructed IDH variants with retained enzymatic activity and possible light-switching capability, we used an *in vivo* screening approach. Specifically, we grew *Saccharomyces cerevisiae* wildtype (WT), and *S. cerevisiae* strains carrying the engineered IDH variants as well as the single and double deletion mutants of IDH1 and IDH2 as spots on two agar plates with yeast extract peptone (YP) medium and 1% acetate as a carbon source. On this carbon source, both IDH1 and IDH2 genes were found essential as single knock-out or as double knock-out^32,33^. We exposed one plate to light and kept a second one in the dark.

To illuminate the strain library spotted on agar plates with a defined amount of blue light, we constructed a device with 96 blue LEDs, arranged in a format that is compatible with 96-well plates (Fig. S2). The device can tune the light intensity of each LED individually over a wide intensity range, while it is also designed to efficiently remove the heat generated by the LEDs. With this device, we screened our library of IDH variants and the control strains. We used the WT, i.e., the parental strain of the library, as a control for potential light toxicity effects, and the single and double deletion mutants of IDH1 and IDH2 as negative controls. The cells were grown on the plate for ~65 hours either with or without light. After this period of time, images were taken with the purpose to quantify the cell growth of each spot (Fig. 2A, Fig. S3).

**Fig. 2:**
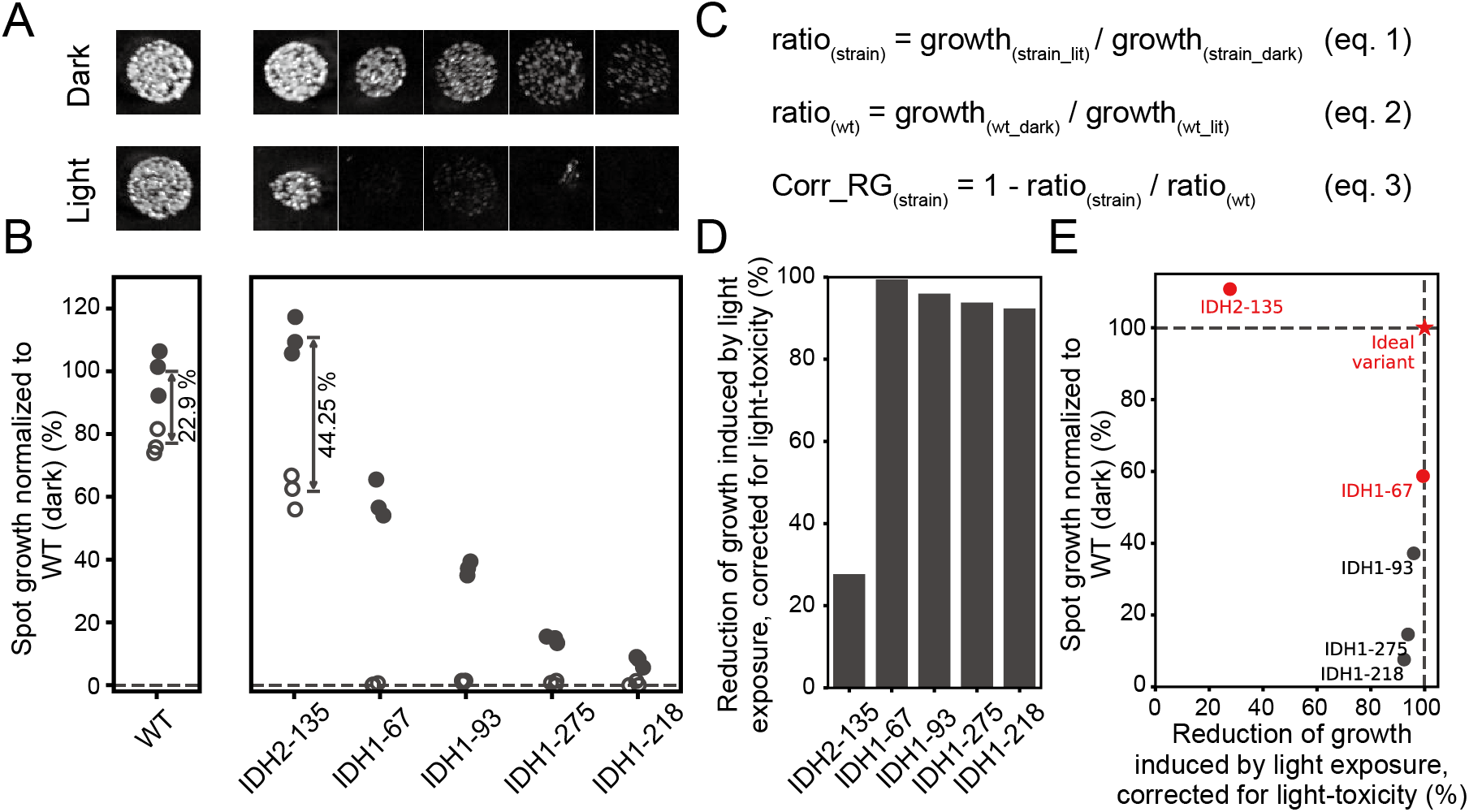
Library screening with spotting assay on 1% acetate. (A) Background-corrected and contrast-adjusted images of representative spots grown on 1% acetate YP medium in dark and light-exposed condition (constantly exposed to light at 451 nm, with an intensity of ~220 μW/cm^2^) from the WT and the five variant strains that had shown growth in the dark condition. Spots of each column correspond to the x-axis of (B). For full image, refer to Fig. S3. (B) Spot growth quantified by total gray scale of the segmented spots, normalized by the mean of WT spots from the dark condition. The arrows and texts show the percentage growth drop between the mean growth of the dark and light-exposed conditions. (C) Equations that were used to determine the reduction of growth due to the photo-switchability of the five IDH variants, after correcting for the photo-toxicity effects. The growth values in eq. 1 and eq. 2 were the means of spots growth of the indicated strain/condition. (D) Results of calculation in (C) were plotted for the five strains. (E) Mean of normalized spot growth in the dark versus corrected reduction of growth were plotted for each of the five strains. The red color indicates the ideal variant and the two variants selected for further study. In the later stage of this study, we found that three non-synonymous mutations were accidentally introduced into the IDH genes during library construction, in both strain IDH2-135 and IDH1-185 used in this screening (see result of IDH1-185 in Fig. S3). Corresponding variants without the three non-synonymous mutations showed similar results (Fig. S5).

Here, consistent with earlier reports^32,33^ and showing their suitability as negative controls, we found that the IDH deletion mutants cannot grow on acetate (Fig. S3). Second, focusing on the experiments in the dark, we found that the WT and five variants grew, but eight variants did not (Fig. S3), suggesting that in these variants the engineered IDH versions are inactive. Third, comparing the WT growth between the dark- and light-exposed conditions, we found that the WT had a lower growth rate under the light-exposed condition (Fig. 2A), which is likely due to the fact that blue light is toxic for cells under respiratory conditions^34^. Similarly, the five IDH variants, which grew in the dark, also grew slower under the light exposure (Fig. 2A).

To quantitatively assess the growth of each variant under the two conditions, we segmented each spot to determine the mean gray scale and normalized all gray scale values to the mean value of the WT in the dark (Fig. 2B). Here, we found that in the dark the IDH2-135 variant grows similarly to the WT, suggesting an fully active IDH when not exposed to light. The other four variants showed somewhat compromised growth in the dark, suggesting that in these variants the IDH structure is affected by the integration of the LOV2-domain (Fig. 2B). For each variant, the differences between dark and light-exposed conditions (Fig. 2B) are a combination of light-toxicity and photo-switching, and therefore we needed to decouple the two effects.

To derive the light-switching effect without the light-toxicity effect, we determined the ratio between the light-exposed growth and dark growth for each strain, as well as the WT (Fig. 2C, eq. 1 and 2). We then corrected for light toxicity effect by dividing the strains’ ratios by the WT ratio and subtracting this ratio from 1 (Fig. 2C, eq. 3), resulting in a measure of the reduction of growth caused by photo-switchability of the engineered IDH. When applying this to the different variants, we found that all five strains with the IDH variants had a reduction of growth that went beyond the light-toxicity effect, suggesting that light-induced conformational changes in the LOV2-IDH proteins have led to reduced IDH activity and thus growth (Fig. 2D).

An ideal, photo-switchable IDH variant would be one that had a normal growth rate in the dark as well as a high reduction in growth rate upon light exposure. Such variant would be situated in the upper right corner in Fig. 2E. Thus, for our further assessment of the different variants, we selected those two variants that came closest to this ideal, i.e., the variant with normal growth rate in the dark but a lower response to light exposure (IDH2-135), and the variant with a high reduction in growth rate upon light exposure (after correction for phototoxicity) but some reduction in growth rate in the dark (IDH1-67). LOV2 insertion at the IDH2-135 site is expected to modulate the catalytic site of IDH2 via an anti-parallel stretch, while insertion at the IDH1-67 site allows direct modulation of an allosteric control region.

To further confirm that insertion of the LOV2 domain at the IDH2-135 or the IDH1-67 sites can achieve photo-switchability of the IDH enzyme without complication of light toxicity, we used LOV2 mutants that are constantly in the dark conformation (LOV2-dark) or in the light-switched conformation (LOV2-lit)^21^. For both the IDH2-135 and the IDH1-67 sites, strains with the LOV2-dark domain inserted have a higher growth rate than the corresponding strains inserted with the LOV2-lit domain (Fig. S4). This demonstrates that the LOV2 domain in the light-activated conformation when inserted at either of the two sites leads to reduced activity of the IDH.

### Functional validation of identified variants

Because our screening experiments only assessed the long-term growth phenotype of a plated yeast cell population, we next sought to further validate the photo-switchability of the two selected IDH variants. Specifically, we aimed to perform dynamic perturbation experiments, with a direct assessment of metabolic phenotypes after turning on the light. First, with the IDH being an enzyme of the CO_2_-producing tricarboxylic acid cycle, we aimed to dynamically measure the carbon dioxide transfer rate in shake flask cultures upon light exposure. However, before we did so, we first estimated the normal (i.e., unperturbed) contribution of the IDH on the cellular CO_2_ production rate, when cells grow in steady-state on acetate. To this end, we used a combined stoichiometric/thermodynamic model and ran flux balance analysis simulations for growth on acetate, where the model was constrained by the earlier identified upper limited in the Gibbs energy dissipation rate^35^.

These simulations identified three major CO_2_ producing reactions: the phosphoenolpyruvate carboxykinase (PEPCK), mitochondrial α-ketoglutarate dehydrogenase (AKGDm), and isocitrate dehydrogenase (ICDH) each contributing about 30% to the overall CO_2_ production rate (Fig. S6). As inhibition of the ICDH reaction will also lead to a halting of the directly following and also CO_2_ producing AKGDm reaction, in principle, upon a sudden inhibition of the IDH, we could expect a drop in the CO_2_ production rate of about 60% (i.e., 30% by IDH, 30% by AKGDm) within minutes, considering typical metabolite turnover rates. However, this estimation disregards flux re-distributions upon IDH inhibition, e.g., from the IDH substrate isocitrate into the glyoxylate shunt to directly feed PEPCK, the third major CO_2_ producing reaction, as well as the malic enzyme, which could continue producing CO_2_ as long as ATP is still available (Fig. S6). Thus, we could maximally expect an about 60% reduction in the CO_2_ production within minutes, but due to flux rerouting, the reduction could also be much less.

To test for reductions in the CO_2_ production rate upon light activation of the engineered IDH, we used a device to monitor carbon dioxide transfer rates (CTR) in shake flask cultures (Fig. 3A). Specifically, we grew yeast flask cultures (wildtype, and the two selected variants with engineered IDH) to exponential phase with minimal medium and acetate as the carbon source. Here, we observed that growth rates for the WT and the strains with the IDH variants, as estimated from the CTR measurements, are consistent with the dark growth rate determined from the spotting assay (Fig. 3B), cross-validating our finding on the basal (i.e., dark) IDH activity in the different strains.

**Fig. 3:**
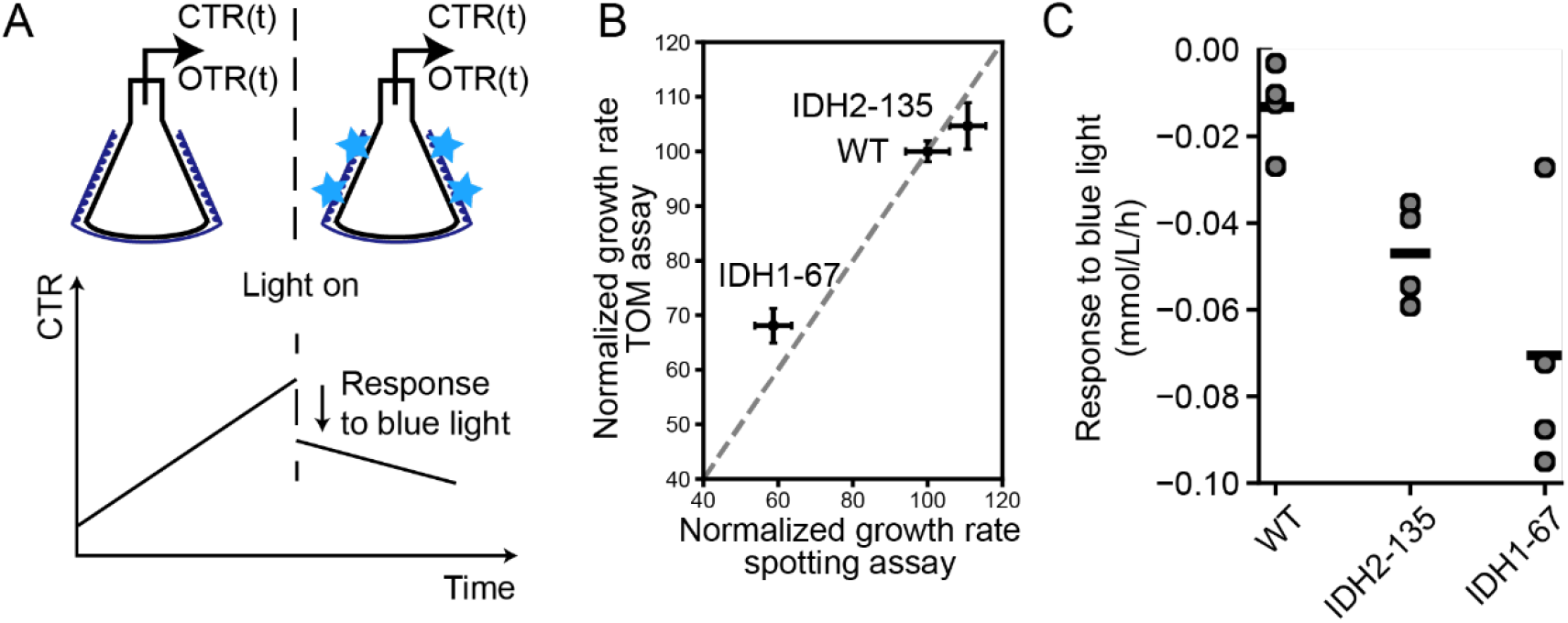
Carbon dioxide transfer rate responses to light perturbation of selected variants. (A) Schematic representation of assays. Flasks were connected to sensors so that CTR and OTR can be measured over time. They were also wrapped with LED strips so that blue light can be illuminated on the cell culture at desired time point. To quantify the responses to light, the first drop of CTR after turning on the light was used. (B). Comparisons of growth rates measured by TOM assay and by spotting assay, both normalized to the mean growth rate of wildtype strain grown under the same condition. Standard deviation and mean from three spotting assay observations or six TOM assay observations were plotted. (C). Responses to light perturbation of wildtype and photo-switchable strains. Results of four biological replicates are shown. The horizontal line of each strain shows the mean of all four measurements. See also Fig. S8.

At this condition of steady growth, we then turned on the light source assembled around the flask (Fig. S7), while measuring the CTR of the culture every 30 minutes. With the WT and the variants, upon light exposure and after an initial increase of CTR likely due to a temperature increase (Fig. S7D, E), a gradual change in the CTR occurred, slowly declining over the course of several hours (Fig. S8A), again indicating a toxic effect of the light. However, we found a stronger drop in the CTR in the strains with the IDH variants between the first and the second measurements after turning on the light compared to the WT (Fig. 3A, Fig. 3C). This further supported that there is indeed a light-switching effect of the engineered IDH enzyme, also upon dynamic perturbation, while the long term drop in CTR is dominated by photo-toxicity (Fig. S8B, C).

### Optimization of variants

Next, we aimed to optimize our light switchable IDH variants with the aim to switch the IDH with less light, and to thereby reduce phototoxicity. To this end, we used LOV2 variants with slow-reverting kinetics, which once activated by blue light will remain switched for several minutes^16^. These LOV2 variants have a single amino acid mutation on the N-helix (V416I or V416L). We introduced these LOV2 variants at the two sites identified in our screen (i.e., IDH2-135 and IDH1-67), obtaining four new strains with presumably slow-reverting kinetics. Here, the logic is that once light-activated, the optogenetic domain will stay longer in this conformation, and we thus would need less light to keep the IDH photo-switched.

First, we again performed spotting assays with these strains, the WT and the original variants. Without light, the slow-reverting LOV2 variants of IDH2-135 had a similar growth rate (normalized to the WT) as the IDH2-135 variant with natural LOV2 domain and the WT strain (Fig. 4A). However, we found that the slow-reverting LOV2 variants of the IDH1-67 strains had a higher growth rates than the natural LOV2 variant of IDH1-67, comparable to the growth rate WT strain (Fig. 4A). We confirmed this finding by growth rate measurements in liquid cultures (Fig. S9). This suggested that the slow-reverting variants of IDH1-67 have higher basal IDH enzyme activity compared to the natural LOV2 variants. Since the three IDH1-67 strains only differ in LOV2 conformation, it further suggests that the IDH enzyme activity is highly sensitive to structural modulation at the IDH1-67 site, revealing good photo-switching potential of the site.

**Fig. 4:**
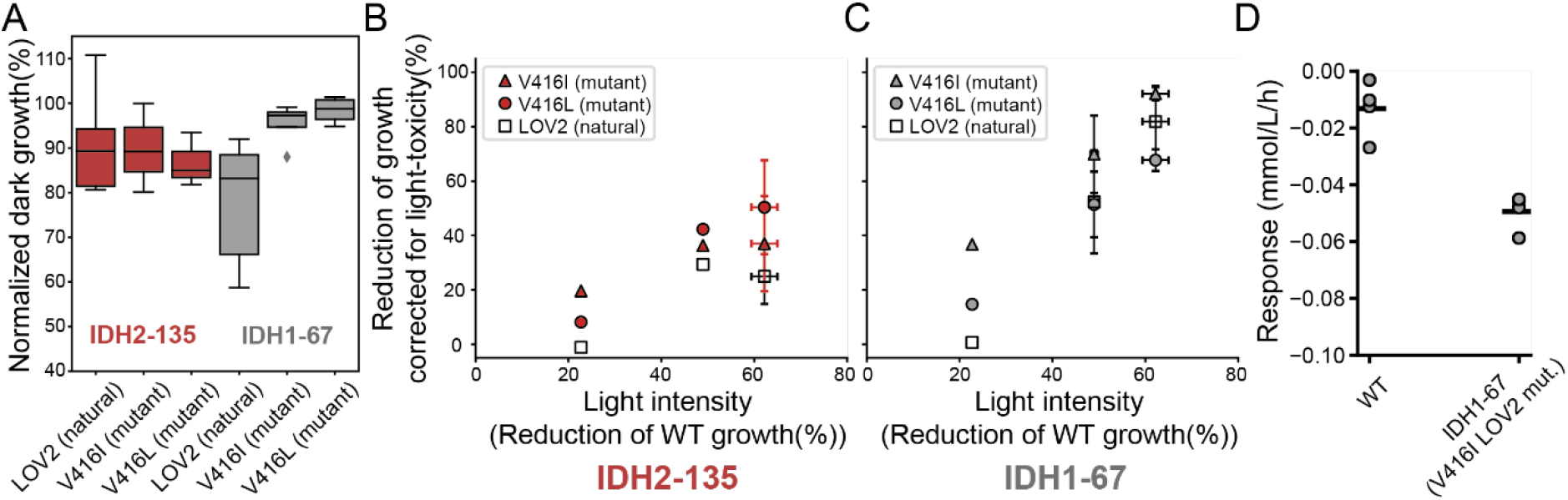
Optimization of variants. (A) Growth of spots in the dark from 3 or more experiments, normalized to the WT strain used in the same experiment. The grey boxes are from IDH1-67 strains while the red boxes are from IDH2-135 strains. (B, C). Response to blue light illumination (reduction of growth induced by light calculated as shown in Fig. 2B), across different light intensities (indicated by the reduction of growth of wildtype spots) for IDH2-135 strains (B) and IDH1-67 strains (C). Error bars (on both the x- and y- axis) for dots from (B) at light toxicity ~49% showed standard deviation from 3 experiments, while error bars (on both the x- and y- axis) for dots from (B, C) at light toxicity ~62% showed standard deviation from 2 experiments. The rest of the dots show results from one experiment. (D) Response of CTR to blue light calculated as in Fig. 3A. The wildtype data here is the same dataset as in Fig. 3A. Data from 4 biological replicates, performed on 2 different days. The horizontal line of each strain shows the mean of all 4 measurements.

For illumination of the different variants in the spotting assays, different light intensities were applied to examine whether we can switch the slow-reverting variants with less light. Here, we used the light-induced reduction of growth of the wildtype as an indication of the light intensity applied in each experiment. As the light intensity increases, all strains showed an increasing response in terms of corrected reduction in growth (defined as in Fig. 2C, D). At all light intensities, the four variants with the slow-reverting LOV2 consistently showed a stronger response than the corresponding variants with the natural LOV2, with the IDH1-67 (V416I mutant) strain having the strongest response at all light intensities (Fig. 4B, C, Supplementary Table S2). This suggests that with this LOV2 variant less light is required to achieve a certain reduction of IDH enzyme activity. We also tested the dynamic response of the IDH1-67 (V416I mutant) strain in flask culture with CTR measurement, where we found that it had comparable photo-switchability as the IDH1-67 with original LOV2 domain but not larger (p = 0.22, T-test, Fig. 4D, Fig. 3C). The fact that the CTR response is not larger for the V416I mutant, as contrary to the growth response during spotting assay could be due to differences in dynamic measurements and long-term growth measurements. Overall, we now had improved photo-switchable IDH variants that have a normal basal activity (since the strains showed a normal growth rate) and better switchability with less light, partly mitigating the problem of light toxicity.

### Application in single cells

To test the performance of our photo-switchable IDH enzyme in single cells, we cultured cells in microfluidic devices with acetate as the carbon source and tested different single-cell readouts. Using UV excitation, we monitored NAD(P)H levels via the autofluorescence of these molecules^3^, as this provides an almost online metabolic concentration readout of important metabolites. Since the IDH enzyme and the TCA cycle generates the majority of NADH during respiration, we asked whether inhibiting IDH activity with our photo-switchable enzyme would cause a drop in NAD(P)H fluorescence. Single cells from different fields of view were either constantly grown in the dark (hence the dark cells), or were first grown in the dark to allow adaptation and then exposed to blue light throughout the rest of the movie (hence illuminated cells). Two sets of experiments were performed with a light intensity of either 4 mW/cm^2^ (Fig. 5) or 40 mW/cm^2^ (Fig. S12).

**Fig. 5:**
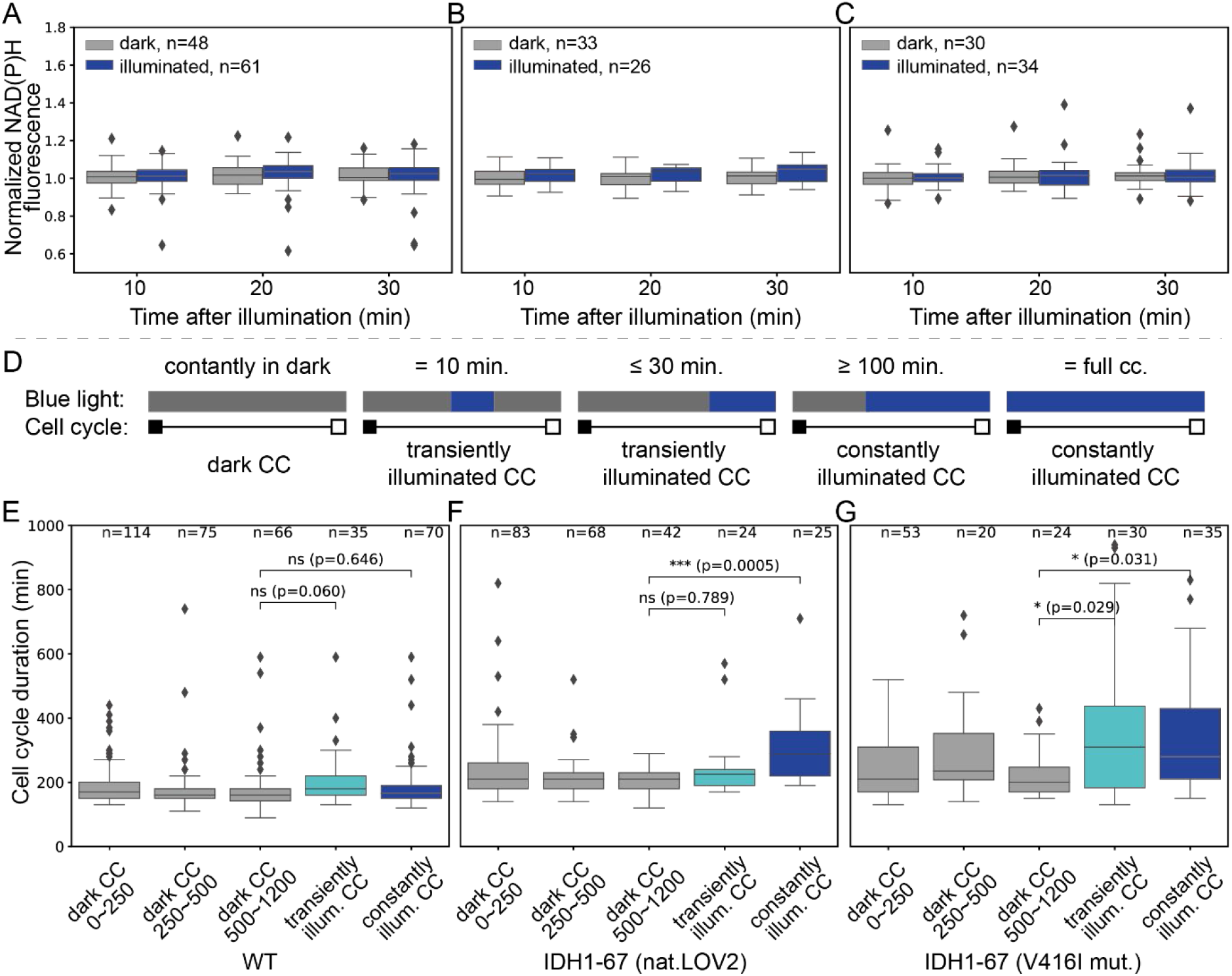
Application in single cells. (A, B, C) Normalized NAD(P)H fluorescence of dark and illuminated cells, of WT (A), IDH1-67 (natural LOV2) (B) and IDH1-67 (V416I mutant) strains (C), collected at three time points after blue light at an intensity of 4 mW/cm^2^ was turned on. To normalize, the NAD(P)H fluorescence at these time points was divided by that of the time point before turning on the light for each single cell. (D). Cell cycles in each experiments were divided into three types: dark CC: CCs that remain constantly in the dark; transiently illuminated CC: CCs that were either illuminated for ten minutes (from the two transiently illuminated fields of view), or were illuminated for 30 minutes or less (part of the CCs that were already in late progress when light was turned on for the constantly illuminated field of view); constantly illuminated CC: CCs that were either illuminated for 100 minutes or more (part of the CCs that were in early progress when light was turned on for the constantly illuminated field of view) or CCs that were illuminated for the full cycle (CCs that started after turning on the light for the constantly illuminated field of view). (E, F, G) Duration of the dark CCs (started at 0-250, 250-500 or 500-1200 minutes of the movie), transiently illuminated CCs and constantly illuminated CCs, of WT (E), IDH1-67 (natural LOV2) (F) and IDH1-67 (V416I mutant) strains (G). (Numbers of CCs plotted for each box are shown on top. p-values of Kolmogorov-Smirnov tests between the indicated data groups are shown, and annotated with * if p ≤ 0.05 or *** if p ≤ 0.001.

Here, we first examined the NAD(P)H response to blue light illumination at 4 mW/cm^2^, an intensity at the same scale as that used for the carbon dioxide transfer rate assay (Fig. S7C). To quantify the response, the NAD(P)H fluorescence of the three time points after turning on the light were collected for each single cell, and normalized to the fluorescence of the same cell at the time point right before illumination (Fig. 5A, B, C). Here we observed that for both the dark and the illuminated cells and all three strains, normalized NAD(P)H fluorescence remained constantly at 1.0, thus 4 mW/cm^2^ blue light illumination did not caused a response in the NAD(P)H fluorescence. We further examined the effect of 40 mW/cm^2^ blue light perturbation. Here we observed that for the illuminated cells, the normalized NAD(P)H fluorescence rose to around 1.1, higher than that of the dark cells at same time points or the fluorescence of the same cells before illumination (which is used for normalization and thus is 1.0) (Fig. S12A, B). Here, the same behavior is observed in both the WT and the IDH1-67 photo-switchable strain, suggesting that this response in NAD(P)H is not due to photo-switching of IDH. This observation could be caused by strong blue light excitation, which could be very toxic for the cells leading to a strong cellular response and potentially masking the effect of IDH photo-switching. Besides, we recently learned that the measured cellular autofluorescence of both NADH and NADPH might stem mostly from NADPH whose concentration is not directly perturbed upon IDH inhibition, as cellular NADPH levels are 100-1000 fold higher than NADH levels (personal communication Martin Pabst, TU Delft). We therefore concluded that NAD(P)H fluorescence is not a good readout to monitor switching in the IDH activity.

We next checked the duration of cell cycles (CC, defined as the time between one budding event to the next budding event) of our light-switchable strains when 4 mW/cm^2^ light was turned on. Yet, as we compare CC durations in the dark and with CC durations when the light was turned on, we first have to examine whether cell aging was a factor affecting the CC durations. To check this, we first divided the CC of cells kept in the dark into three groups. Here, we found that cell aging does not affect the CC duration, as CC that started at early (0~250 minutes), in the middle (250~500 minutes) and late (500~1200 minutes) during the experiment have similar duration in all three strains (Fig. 5 E, F, G, grey boxes).

We further asked whether the CC perturbed by light were significantly longer than the CC in the dark for each strain. Here we also investigated the effect of different illumination time, by turning on the light at two fields of view for only ten minutes, while keeping the light on for a third field of view constantly for the rest of the movie. Together with the dark field of views, in total we analyzed the following types of CC: dark CC, transiently illuminated CC, constantly illuminated CC (Fig. 5D). Since blue light was turned on at around 500 minutes for all three strains, we compared the dark (500~1200 minutes) group to the constantly illuminated group and the transiently illuminated. Here, we found that for the WT strain, blue light illumination did not change the CC duration (Fig. 5E), when analyzed by a Kolmogorov-Smirnov test. However, blue light illumination prolonged the CC duration of the constantly illuminated CC for the variant IDH1-67 (natural LOV2) (p < 0.001, Fig. 5F), as well as the CC duration of the both the transiently illuminated CC and constantly illuminated CC for the variant IDH1-67 (V416I mut.) (p < 0.05, Fig. 5G). This confirms that the engineered IDH is also switchable when perturbed in single-cell level.

Remarkably, we did not observe clear phototoxicity at the light intensity of 4 mW/cm^2^ in the microfluidic experiment, for the WT cell, while we observe clear toxicity effect in the spotting assay and the CTR measurement assay. For the spotting assay, although we were using an even lower light intensity (220 μW/cm^2^, Fig. S2D), the cells were grown for a much longer period (~65 hour) and condition could be more stressful on agar plate. For the CTR measurement assay, however, the timescale and the growth condition were similar. The phototoxicity there might be explained by two factors: (i) cells grown in flask liquid culture might receive more illumination than measured. With the setting of the power source, we measured an intensity of around 1.5 mW/cm^2^ (Fig. S7C) with a sensor. Since the sensor is measuring light shedding on a plane from one side, while the cells were receiving light from all direction, the actual amount of light received by the cell could be maximally four times higher than measured, as inferred by the ratio of the surface area of a sphere to the area of its cross-section plane, making it 6 mW/cm^2^, higher than that used in the microfluidic experiment. (ii) Cells in the CTR measurement assay also experienced a rise in temperature, at least by 1.5 degrees Celsius as measured in the liquid culture by the end of the experiment, but could also be higher (Fig. S7E). This heat came from the dissipation of the LED strips, and it could have stressed the cell alongside the light toxicity.

## Discussion

In this study, we rationally designed the first version of a photo-switchable IDH enzyme in yeast, for which we exploited its allosteric regulation mechanism. We accomplished this by first computationally identifying suitable insertion sites for the optogenetic domain LOV2 and subsequently *in vivo* screening of variants using a simple growth assay, where IDH activity is essential. While phototoxicity caused by the light exposure is still an issue, we showed in growth assays on agar plates, assays where we monitored the CO_2_ transfer rates in flask culture, as well as single cell assays where we monitored the cell cycle duration that the activity of the engineered enzyme is indeed modulated by light. While further optimization of the IDH variants is still needed, our work showed that novel metabolism perturbation tools could be developed by inserting optogenetic domains into rate-limiting enzymes.

Unlike previous studies that engineered photo-switchable enzymes *in vitro*^23,24^, we screened directly *in vivo* for enzyme variants that could be switched by light, using a screening scheme with yeast growth as output parameter. We opted for this route due to the discrepancy of protein function and biochemical properties *in vivo* and *in vitro*^36^. However, it turned out that the detection of *in vivo* photo-switchability and its quantitation can be obscured by multiple factors, such as the re-distribution of metabolic flux and potentially silent (i.e., growth) phenotypes, and non-linear relationship between enzyme activity and readout. Therefore, rapid and direct readout of the substrate and product of target enzyme using metabolomics methods with fast sampling would be the ideal and generalizable method during screening and for characterization of variants.

*In vivo* application of optogenetics also involves potential phototoxicity on cells, which is not only unwanted, but also complicates quantification of the photo-switchability, as seen in this study. In fact, the unwanted effect of the light on cells is still a major issue of the here developed photo-switchable IDH enzyme. In this study, we only used respiratory conditions to test the sensor. Under these conditions, we expect phototoxicity to be more severe compared to fermentative conditions, as light-toxicity has been shown to mainly affect the respiratory chain^34^. For future application of the engineered IDH variant, it is necessary to reduce photo-toxicity. This could be done by (i) increasing the effect of the light-induced changes in the LOV2 domain onto the IDH structure or by (ii) using a LOV2 domain that can be activated by less harmful light. For the first point, we have already shown that slow-reverting LOV2 variants have a positive effect. For the second point, LOV2 variants that can be switched by light at a different, i.e., higher, wavelength could be used. In fact, efforts in this direction are ongoing; colleagues have recently identified LOV2 variants that can be activated at 520 and 620 nm. Unfortunately, however, these variants require enormously high light doses, meaning that they are not yet suitable for *in vivo* applications. Further, by deleting the NADP+ dependent IDH enzyme (IDP1/IDP2/IDP3), NAD^+^-dependent IDH enzyme might also become essential under fermenting conditions, and could therefore allow for photo-switching test with lower light toxicity.

The different growth rates of the IDH1-67 variants with the different LOV2 mutants suggested that IDH enzyme is highly sensitive to structural modulation at the IDH1-67 site (Fig. S9). Therefore, further improvements can be envisioned by adding various additional linkers between the LOV2 domain and the IDH enzyme at this site, which might reduce the structural hindrance of the LOV2 domain on the IDH enzyme and improve its basal activity when LOV2 domain is not activated by light. On the other hand, both the IDH1-67 and the IDH2-135 sites are embedded in surface loops containing three residues (Table S1), so truncating one or two of these residues might further improve photo-switchability of the IDH variants, as that could aid transmission of structural change in the LOV2 domain to the IDH enzyme. This is of particular interest for the IDH2-135 variant as it already has a normal basal activity (Fig. 2B), but is also of interest for IDH1-67 variant and might be combined with linker adding.

## Supporting information

Supplemental items

## Acknowledgement

Funding from the following sources is acknowledged: the Chinese Scholarship Council (for H.C.). H.C. would like to thank Marco Fraaije and Friso Aalbers for helpful discussion of library design and characterization, Wigger Jonker and Harry van Driel for help with electronics and mechanical work.

## Author contribution

Conceptualization: H.C. and M.H.; Library design: H.C. and H.W.; Library construction and screening: H.C., L.M.; Optimization and further characterization: H.C., R.W., J.P.V.C.L., M.R.; LED device programming: T.C.L.; Formal analysis: H.C., L.M. and A.P.; Writing-original draft: H.C., H.W. and M.H.; Writing-review and editing: H.C., M.H.; Supervision: M.H.; Funding acquisition: M.H. and H.C.; Project administration: M.H.

## Materials and method

### Computational identification of insertion points

To identify insertion points for the LOV2 domain, the method of the Hahn group^21^ was used with some modifications. A script written in yasara^27^ was used to calculate the surface accessible area, the secondary structure, and the Cα-Cα contact distances from the pdb file 3blx^25^. Kullback-Leibner conservation scores were calculated by MISTIC^37^ using alignments obtained from the PFAM database^38^. To calculated the Cα-Cα distance between any two residues, we considered all four copies of these residues within the hetero-octamer. That is, for each two residues, 16 Cα-Cα distances can be calculated. Residues were considered to be contacting if their shortest Cα-Cα distance (out of the 16) was less than 7 Å. The allosteric control regions of IDH were described by Taylor *et al.*^26^ while the catalytic residues were those that lie within the active site of IDH^25^ (Fig. 1A).

### Yeast strains construction

Yeast strains used in this study are based on the IMX585 strain^31^, which is prototrophic and of the CEN.PK113-7D strain-background. A *Cas9* gene under the control of the *TEF1* promotor was integrated into the *CAN1* locus together with the *natNT2* marker. To knocked-in the LOV2 domain at the 13 selected insertion sites (Table S1) with *Cas9*, primers containing the 20 bp spacerthat target the nearby sequence of each insertion site were designed with aid of the Yeastriction webtool^31^, so that double strand break (DSB) can be created near the insertion sites. Among the 13 insertion sites some were relatively close and thus could be targeted with the same sgRNA. In total, eight sgRNA sequences were selected and integrated into the expression plasmid pROS13A. Repair fragments consisting of 60 bp homologous arms that flanked both sides of the LOV2 domain or LOV2 mutants were generated by PCR using template plasmids containing the LOV2 sequences. The 60bp homologous arms determined the exact insertion sites, covered the sgRNA-targeted cutting site, and contained 3 synonymous mutations on the sgRNA pairing sequence to prevent recurrent cutting of Cas9 after DNA integration.

For the construction of dark and lit variants, LOV2_I510E_I539E_ (lit-mutant) and LOV2_C450M_ (dark-mutant) were synthesized chemically and used for the generation of repair fragments. The same primers and sgRNA expression plasmids were used as for the generation of the corresponding strain with natural LOV2 domain. For the construction of the repair fragments of the slow-reverting strains, template plasmids with LOV2 sequence carrying the V416I or the V416L mutation were created via a two-step PCR-directed mutagenesis, where the template plasmid was first PCR amplified into two overlapping fragments using primers that contain the desired mutation, and the assembled via Gibson Assembly^39^ and validated by Sanger sequencing. For the construction of control strains with deleted IDH genes, three distinct plasmids that targeted either *IDH1*, *IDH2* genes or both were constructed. Repair fragments for deletion were chemically synthesized and annealed. For an overview of all primers and plasmids refer to Table S3 and Table S4, respectively.

For yeast transformation, single yeast colonies growing on yeast extract-peptone-dextrose (YPD) 20 gL^−1^ glucose agar plates were used to inoculate 10 mL YPD medium in 100 mL culture flasks, and grown at 30°C, 300 rpm. The transformation reactions were performed as described previously^31,40^. After obtaining single colonies on selective YPD plates, single colonies were picked and re-streaked on empty YPD plates, after which the newly grown single colonies were picked and restreaked again, to ensure homogeneity in the genetic background of single colonies. These secondly re-streaked colonies were then inoculated for genome extraction, PCR assay and Sanger sequencing to verify the genetic modification. Frozen yeast stocks were produced by adding 400 μL of glycerol (85%~89%) to 600 μL *of S. cerevisiae* overnight culture in YPD and stored at −70 °C. The strains were revived from the −70 °C freezer on YPD agar plates. For an overview of all strains, refer to Table S2.

### LED device construction

A LED device was constructed to allow simultaneous screening of all IDH-LOV2 variants illuminated at 451 nm (Fig. S2). The printed circuit board of the device contains 96 High Power LEDs (GD CS8PM1.14, OSRAM) that are positioned in alignment with a 96-well plate. Individual control of all 96 LEDs is achieved through 8 TLC5940 16-channel LED drivers (Texas Instruments) connected in series, of which each driver controls one row of 12 LEDs. Each channel of the LED driver has an individually-adjustable 12-bit grayscale (GS) pulse width modulation (PWM) brightness control and a 6-bit constant-current DOT correction (DC) brightness control. The DOT correction allows for subtle correction of LED brightness difference, which could occur even when the LEDs are under the same voltage and pulse width modulation control, due to inherent heterogeneity of certain LEDs products. The GS and DC matrixes of the 96 LEDs, as well as illumination patterns are programmed by an Arduino Uno microcontroller board that is connected to the serial interface port of the TLC5940 LED driver. Notably, DOT correction was not applied in our experiments, as the measured light intensity of LEDs were similar (Fig. S2D). To assess light intensities of a certain experiment, the light sensor S120VC and module PM100USB from Thorlab was used, with wavelength set as 451 nm. A black plastic lid with drilled holes equal to the size of a spot was used to block scattering light on the sample specimen.

The power supply is facilitated by a 5V power adapter. A DC-DC converter is used in parallel with multiple 0.1 μF ceramic capacitors to smooth current fluctuations in the circuit and to have a well-regulated LED supply voltage. The maximum output current of all channels is determined by an external single resistor, R_(IREF)_, which is placed between the reference current terminal pin and the ground pin. The voltage on IREF, V_(IREF)_ has a value of 1.24 V. The maximum channel current is equivalent to the current flowing through R_(IREF)_ multiplied by a factor of 31.5. To accommodate a greater range of grayscale steps at a lower light intensity, as the screening is performed in vivo, a resistor of 649 Ω was chosen. The maximum current per LED is therefore 60 mA.

In addition, the LED device is augmented with accessory components. A fan-assisted aluminium heatsink (Fisher Elektronik) provides thermal control. A milled PVC microplate holder minimizes the crosstalk of light between the wells and can optionally provide supplementary thermal control through air flow if connected to an air pump. The PVC holder and microplate are hold in place by several aluminium elements attached to the heatsink.

### Spotting growth assay

To obtain a culture for spotting assay, single colonies of yeast strains were inoculated in YPD 20 gL^−1^ glucose medium and grown overnight at 30 °C, 300 rpm. The cultures were diluted to an OD600 of 0.25 and grown for 6 hours to an OD600 of around 2.0. The cultures were then diluted to OD_600_ 0.02 using fresh YPD, of which 3 μL was spotted on two Omnitray Single-well plates (Nunc 242811) with 50 mL YP agar containing 10 gL^−1^ acetate (added in the form of sodium acetate trihydrate) and 100 mgL^−1^ ampicillin to prevent bacterial contamination during long incubation. Technical replicates were made by spotting multiple times.

To align the drops of culture to the LEDs array, the plastic lid of the single-well plate (used for spotting assay) was aligned to the LED array and drawn with dots indicating the positions of each LED. This plastic lid fits well under the single-well plate and aid alignment during spotting. The spotted drops of cells were grown for 65 hours at 30 °C where one of the plates was grown in the dark and the other plate was grown on the LED device with indicated illumination setting, and images of each plate were taken with a Fujifilm LAS-3000 gel imager in EPI mode. To gain quantitative representation of the cell growth from each individual spot, the images were inverted and background subtracted in ImageJ with a rolling ball algorithm with a radius of 75 pixels. An elliptical segmentation was performed for each spot in ImageJ. To segment the spots with very low growth, the display contrast was adjusted to enhance visibility (this is not done for quantitative analysis of the spot). The images were subsequently read by a Python2.7 script where spot segmentations were reproduced by using the coordinates measured by ImageJ. A threshold of 1000 was set to filter out un-subtracted background, and grayscales values above the threshold were summed, normalized by the area, and used for further analysis as indicated.

In the spotting assays for the slow-reverting variants with different light intensities, despite effort to align the spots with the LED, in some experiments there was still systematic misalignment that caused consistent shift of the spots away from the LEDs. As the LEDs have a fixed illumination angle, this shift had an effect in the amount of light exposed on cells. To correct for that, the light toxicity on the wildtype cells of each spotting was used as an indication for light intensity, which accounts for both different LED intensity setting and the shift (Figure 4B and 4C).

### Carbon dioxide transfer rate assay

Single yeast colonies were inoculated into 10 mL of YPD with 20 gL^−1^ glucose and incubated for approximately 10 hours to an OD_600_ between 2.0 and 10.0, dependent on the lag phase of the strain. Proper amount of cells were spun down at 3900 rpm 4 minutes, 30 °C and washed one time with 30 mL pre-warmed Yeast-Nitrogen-Base 10 g·L^−1^ Acetate (YNBA). They were then spun down again, re-suspended and diluted in 50 mL pre-warmed YNBA in 500 mL Erlenmeyer flasks, to a final OD_600_ of 0.1. These flasks were then put into the incubator for approximately 72 hours to adapt to the new medium and grow. When the culture reached an OD between 0.4~1.0, cells were diluted with YNBA to 7.5 × 10^5^ cells/ml or less to a final volume of 50 mL, in flasks manufactured for the Kuhner TOM device (TOM flasks). TOM flasks containing cultivation were attached to the shaker tray with sticky mats (Kuhner Shaker SMX833001) and connected to the TOM device which provide the culture with humidified air flow and measures the CO_2_ transfer rate (CTR) for a duty cycle of 30 minutes, in which for 22 minutes air exchange happened freely, and for 8 minutes the air valves were closed to accumulate CO_2_ and allowed measurement.

The cultivations were grown in the shaker at 30 °C, 200 rpm to a CTR of 1.5 mmol/L/h, which is within the exponential phase. LEDs wrapped around the flasks were then turned on. The cultures were grown for another eight to twelve hours before the experiment was stopped. For illumination, blue LED strips (Okaphone Elektronika LB12M130BN) were fixed on cone-shape 3D printed parts (Fig. S7B) that fit the shape of the TOM flasks. To ensure comparable illumination on technical replicates and to avoid the batch-to-batch heterogeneity of LEDs, a LED strip with 150 LEDs was cut into two halves and applied to two flasks. When mounting the TOM flasks, these parts were also mounted and connected to a digitally adjustable power source (Joy-IT JT-DPH5005). The light intensity of the setup had been measured by using a standard photodiode power sensor (Thorlabs S120VC) from inside an illuminated TOM flask (Fig. S7A). For perturbation, a fixed voltage of 10.5 V was provided to the LED strip, yielding an intensity of around 1.5 mW/cm^2^ when measured from one direction using the sensor. Temperature of the culture was measured with a thermometer after stopping the experiment without turning off the light (Fig. S7E). To estimate the time of temperature rising in the LED, current and power values of the LEDs were recorded from the power source panel at different time point after turning on the light, under the assumption that the rise in power was due to temperature change in the LED strip.

For T-test, the two-sided test for the null hypothesis that 2 independent samples have identical average (expected) values was used, with the assumption that the populations have identical variances. The python function scipy.stats.ttest_ind from the scipy package was used.

### Flux balance analysis simulations

Flux balance analysis simulations were performed based on a novel methodology that integrates the thermodynamic information of each reaction (its Gibbs energy of reaction), while enforcing that the appropriate energy and mass balances are obeyed^35^. The thermodynamic/stoichiometric metabolic network for the yeast *Saccharomyces cerevisiae* was used^35^, with 33 additional reactions (total: 287) and 21 additional metabolites (total: 171). This modified network considers biomass to be a conglomerate of different components (DNA, RNA, proteins, lipids, storage and cell-wall polysaccharides), whereas the former did not distinguish between such macromolecular constituents.

A set of thermodynamically consistent standard Gibbs energy of reaction (Δ_r_G^0^’) was obtained by regression of the model to experimental data (reaction fluxes and metabolite concentration) of yeast cultures grown on glucose. The upper and lower bounds for the metabolite concentrations and Gibbs energy of reaction (Δ_r_G) were then obtained by variability analysis, as previously described^35^. Given that the sign of Δ_r_G defines whether a reaction occurs in the forward or backward direction, training the model with glucose meant that growth on acetate was not allowed. Thus, the bounds of Δ_r_G were relaxed (i.e. widened) and flux balance analysis using growth rate maximization as the objective function was performed. Acetate was used as the sole carbon source and its uptake rate was fixed at different values.

### Microscope

Cells were cultured in the same way as described in the carbon dioxide transfer rate assay until the first YNBA culture reached an OD between 0.4~0.7. Then, the culture was diluted again and cultured for another 24 hours (approximately 5 generations) to obtain a fully adapted and exponentially growing cultures. The culture was then diluted to 0.05 using the same medium and loaded to a microfluidic chip after 5 hours, as previously described^8^. A Nikon Ti-E inverted microscope equipped with a Nikon Perfect Focus System, an Andor iXon Ultra 897 EM-CCD, and a CoolLed pE2 or a Lumencor Aura II light excitation system was used. Bright field images were taken with a halogen lamp as the light source, the light of which was passed through an ultraviolet light filter (420-nm beam-splitter). For NAD(P)H measurements, cells were excited with 365 nm (20% intensity, and 200 ms exposure), with a 350/50-nm band-pass filter, a 409-nm beam-splitter, and a 435/40-nm emission filter. To calculate cell cycle duration, budding events of each cell were manually tracked.

To treat cells with blue light illumination, we used a multiwavelength patterned illuminator (Polygon400, Mightex) integrated into the microscope, at a wavelength of 455 nm to illuminate all pixels of a selected field of view. The Polygon illumination shared the light path of DIC channels, with ND filters installed to reduce the light intensity by either 100-(for high intensity light, Fig. 5) or 1000-fold (for low intensity light, Fig. S12). The Polygon is controlled by independent software from the microscope, and is manually switched on at the desired time point. To constantly illuminate a target field of view while keeping other field of views in dark, we kept the Polygon constantly on, and set the target field of view and the DIC channel as the last point in the microscope job queue. To measure the intensity of the Polygon, the light sensor S120VC and module PM100USB from Thorlab was used. Masks of two different sizes were set using the digital mirrors to constrain the light beam to illuminate a measurable area. Twenty data points were measured at each Polygon intensity setting, and the maximum of measurements was used to represent the light intensity at such setting.

For Kolmogorov-Smirnov test on two samples, the python function scipy.stats.ks_2samp from the scipy package is used. The null is that 2 independent samples are drawn from the same continuous distribution, and we used the default “two-sided” hypothesis.

## Notes

### Competing Interest Statement

The authors have declared no competing interest.

