## Supplemental items for "A photo-switchable yeast isocitrate dehydrogenase to control metabolic flux through the citric acid cycle"

### Supplementary Materials

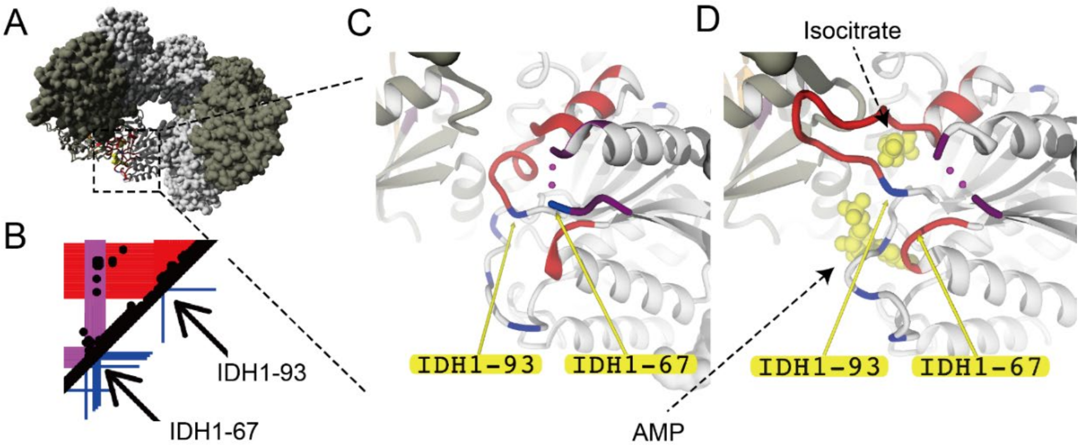


**Fig. S1: Structure and contacts at IDH1-67 and IDH1-93 insertion site.** Related to figure 1. (A). Position IDH1-67 and IDH1-93 are located on the back of the holoenzyme. Color scheme is the same as Fig. 1A. (B). Zoomed-in contact map showing that IDH1-67 is located within an allosterically altered region (pink) and is in contact with the IDH1-93 within the allosteric binding region (red). (C). Local structure of IDH1-67 and IDH1-93 in unbound state. (D). Local structure of IDH1-67 and IDH1-93 with allosteric regulators bound.

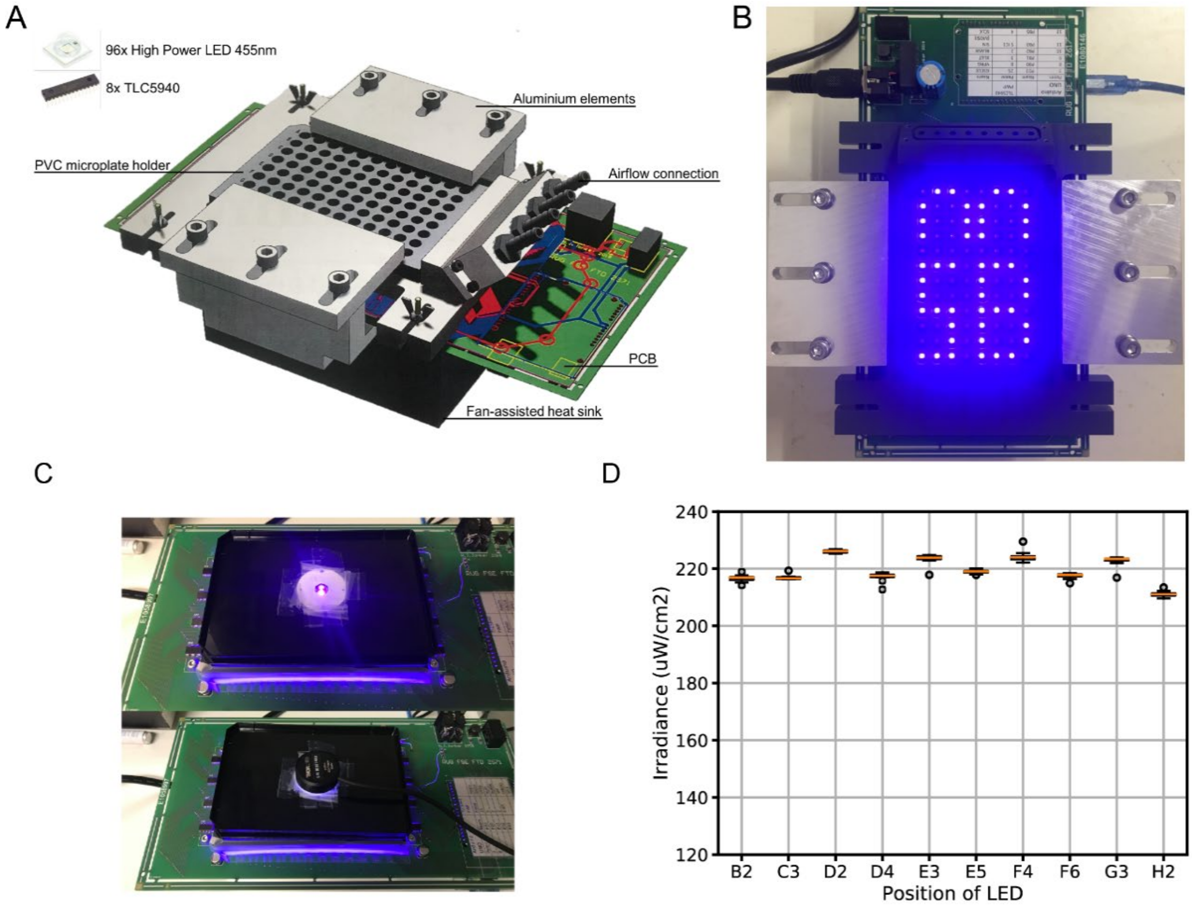


**Fig. S2: LED device and light intensity measurements. Related to Figure 2 and Figure 4.** (A). Design model of the LED device showing different parts. (B). LED device at work. Individual LEDs can be addressed by combination of TLC5940 controller and an Arduino board. (C). Measuring light intensities of an individual LEDs. A black lid with drilled hole equal to the size of a spot was used to block scattering light. A Thorlab S120VC sensor was used for measurement. Measurement was taken at the specimen height. (D). Light intensity of LEDs represented by irradiance ($\mu W/cm^2$). Boxes represents the first and third quantile of 11 measurements and whiskers extend to show 1.5 quantiles.

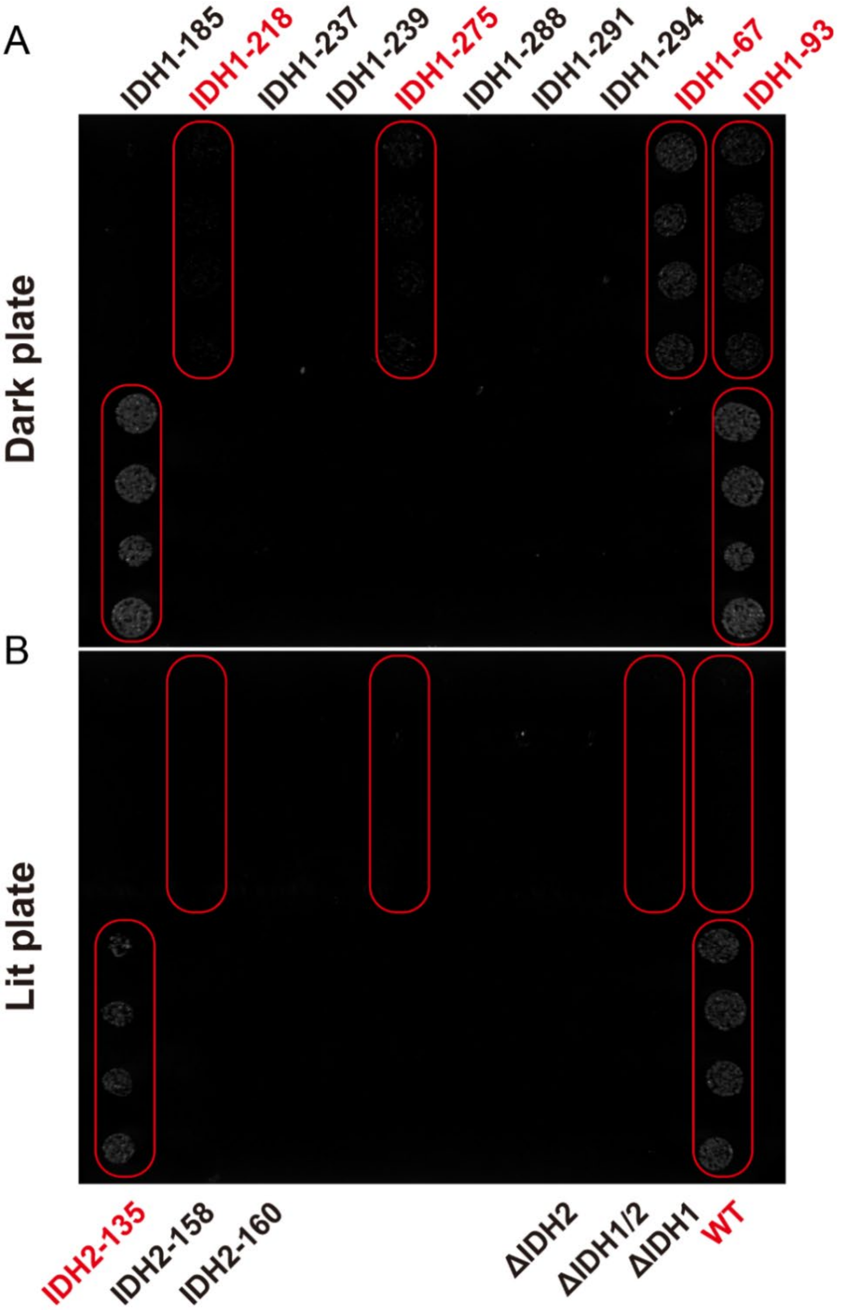


**Fig. S3: Spotting assay for library screening. Related to Figure 2.** Images were taken after 65 hours of incubation at 30 degrees. (A). Agar plate in dark condition. Five out of 13 engineered strains showed visible growth in the dark condition. The IDH knocked-out mutants showed no visible growth. The IDH1-185 and IDH2-135 strains in this experiment contains 3 non-synonymous mutations on the CRISPR seed region, which were introduced by mistake. After correction, the IDH1-185 (corrected) strain showed minor improvement in growth, while the IDH2-135 (corrected) strain showed no difference. (B). Agar plate exposed to blue light during incubation. In general, growth of strains was reduced compared to the dark plate.

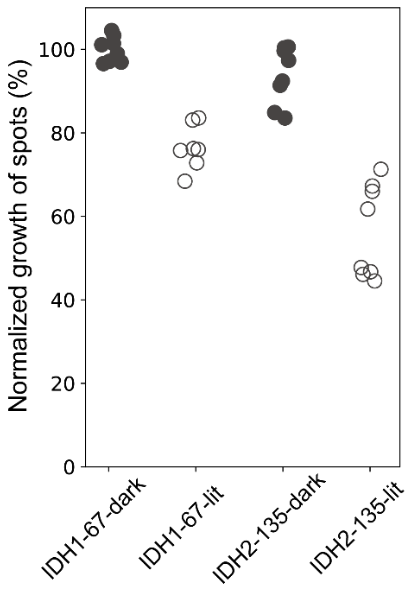


**Fig. S4: Spotting assay of strains with either the dark or lit LOV2 mutant.** Strains inserted with a LOV2 dark mutant (i.e., constantly in the dark conformation) showed a higher growth in the spotting assays than when a LOV2 lit mutant (i.e., constantly in the illuminated conformation) was inserted. Data from two replicates of spotting assays are shown. In each assay, four spots of each strain were grown in the dark at the same condition as in the screening in Fig. 2. For convenience of comparison, the growth of each spot was normalized to the mean of the four IDH1-67-dark spots in each assay. One outlier data point where an excess number of cells were initially spotted was removed for the strain IDH1-67-dark.

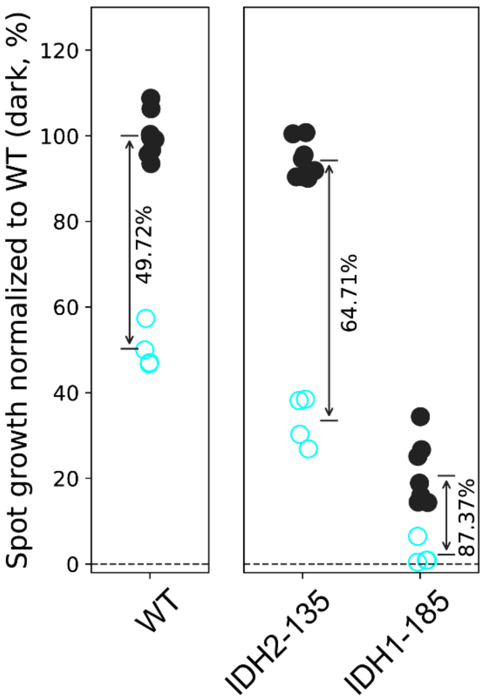


**Fig. S5: Spotting assay for IDH2-135 and IDH1-185 after correction for non-synonymous mutations introduced during cloning.** Spot growth quantified by total gray scale of the segmented spots, normalized by the mean of WT spots from the dark condition. The arrows and texts show the percentage growth drop between the mean growth of the dark and light-exposed conditions. After correction for non-synonymous mutations, the IDH1-185 strain still has impaired (albeit improved) IDH activity that results in slow growth in the dark (Fig. S3), while the IDH2-135 strain remains moderately light-switchable.

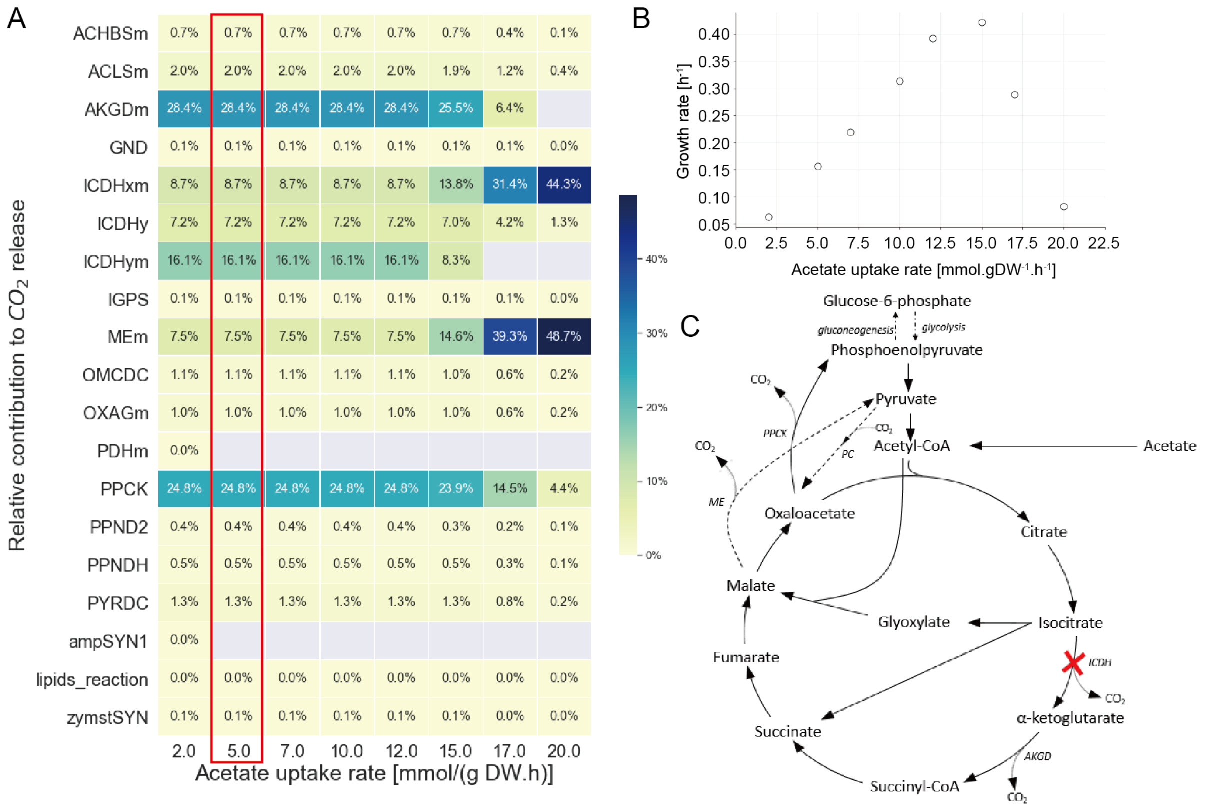


**Fig. S6: Flux balance estimations on $CO_2$ producing reactions. Related to Fig. 3.** (A) The heatmap shows all the reactions contained in the thermodynamic/stoichiometric model that release $CO_2$ (names as used in the model), and their contribution relative to the total $CO_2$ excretion rate. The values are shown as a function of the acetate uptake rate, fixed at different values in the range 2 – 20 mmol/(gDW·h). The relative contribution of the different reactions remains constant (i.e., fluxes scale linearly with the acetate uptake) until an uptake rate of ~15 mmol/(gDW·h), at which the cells are predicted to hit their "upper limit" of Gibbs energy dissipation[1], thereby forcing to a redistribution of metabolic fluxes. Red rectangle marks flux distribution at an acetate uptake rate of 5 mmol/(gDW·h), the condition considered in the main text (see Fig. S6B). Relevant model reactions, mentioned elsewhere in the text: PPCK = phosphoenolpyruvate carboxykinase; AKGDm = mitochondrial α-ketoglutarate dehydrogenase; ICDHxm, ICDHym and ICDHy = isocitrate dehydrogenase. (B) Model predicted growth rate (flux of biomass) on across different acetate uptake rate (AUR). When AUR=5, the model predicts a growth rate that resembles experimentally measured growth rate of a wildtype batch culture (0.154 $h^{-1}$). (C) Schematic representation of the TCA cycle and acetate uptake. Upon inhibition of isocitrate dehydrogenase (ICDH), flux may be redirected via the glyoxylate shunt, eventually leading to an increase in flux (and thus $CO_2$ release) at the level of phosphoenolpyruvate carboxykinase (PPCK) and malic enzyme (ME). At the same time, it is also possible that other reactions - not shown in the diagram above - replenish the pool of α-ketoglutarate, whereby the enzyme α-ketoglutarate dehydrogenase (AKGD) remains active.

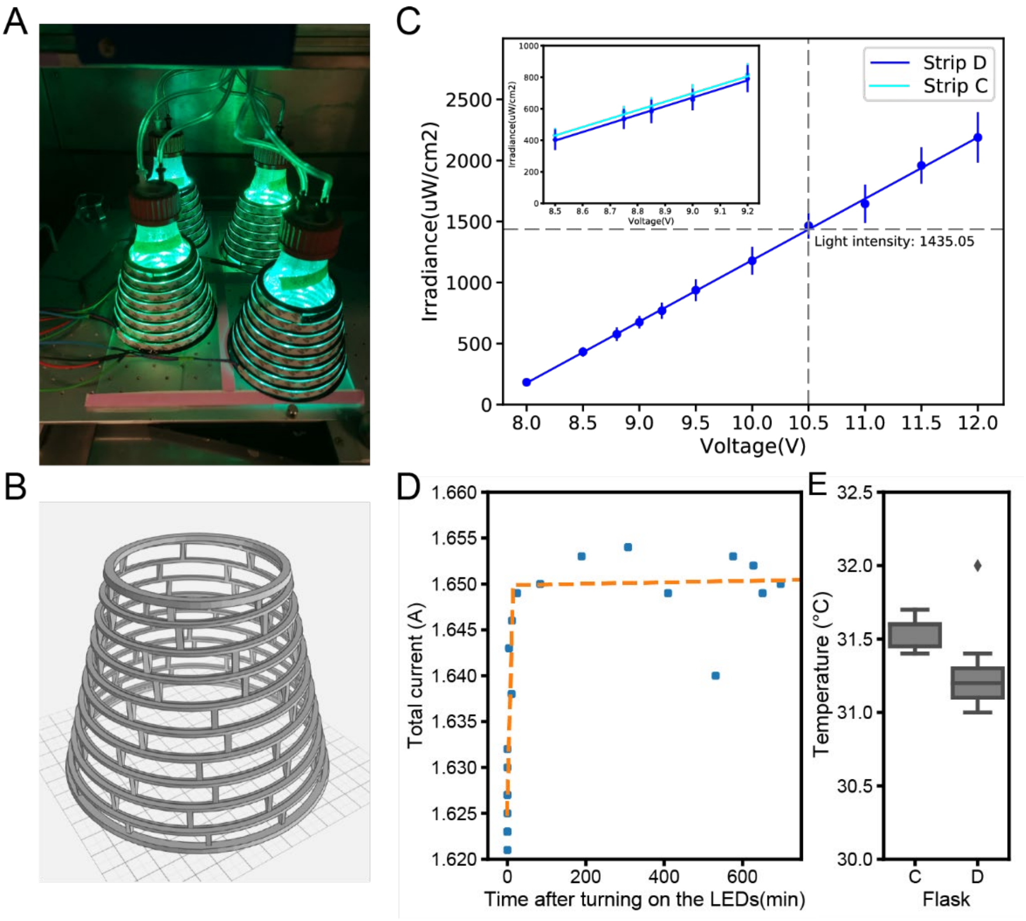


**Fig. S7: Characterization of LED strip setup for flask culture illumination. Related to Fig. 3.** (A) LED strips are fitted in a 3D-printed parts and mounted on flasks. The flasks are connected to the TOM device where carbon dioxide and oxygen exchange rates are measured. Shown are LED with adjustable RGB wavelength. (B). 3D model for LED mounting cone. (C). Light intensity was measured by putting the light sensor S120VC inside an empty TOM flask wrapped with the LED strip and pressed against the glass wall of the flask. The sensor was moved around along the glass wall while 30 data points were taken, the mean of which used as the intensity. The measured intensity, shown as irradiance ($\mu W/cm^2$), is linear correlated with voltage over a certain range. Different LED strips are consistent. Shown here are measurements from two half-strips that were cut from the same strip LB12M130BN. (D). Total current that runs through the system at various time points after turning on the LEDs, recorded by the digital power source from eleven experiments. The orange line is a two-state linear model fitted to the data. The current through the LED strips rise within ~15 minutes (intersection of the two linear model) after turning on the LEDs, presumably due to a rise in the temperature and drop in resistance. (E). Temperature of the cell culture by the end of the experiments. Results from seven experiments were shown for each flask. Confirming our explanation of temperature rising, the temperature of the culture by the end of the experiments were measured to be around 31.5 degrees, instead of 30 degrees (shaker temperature).

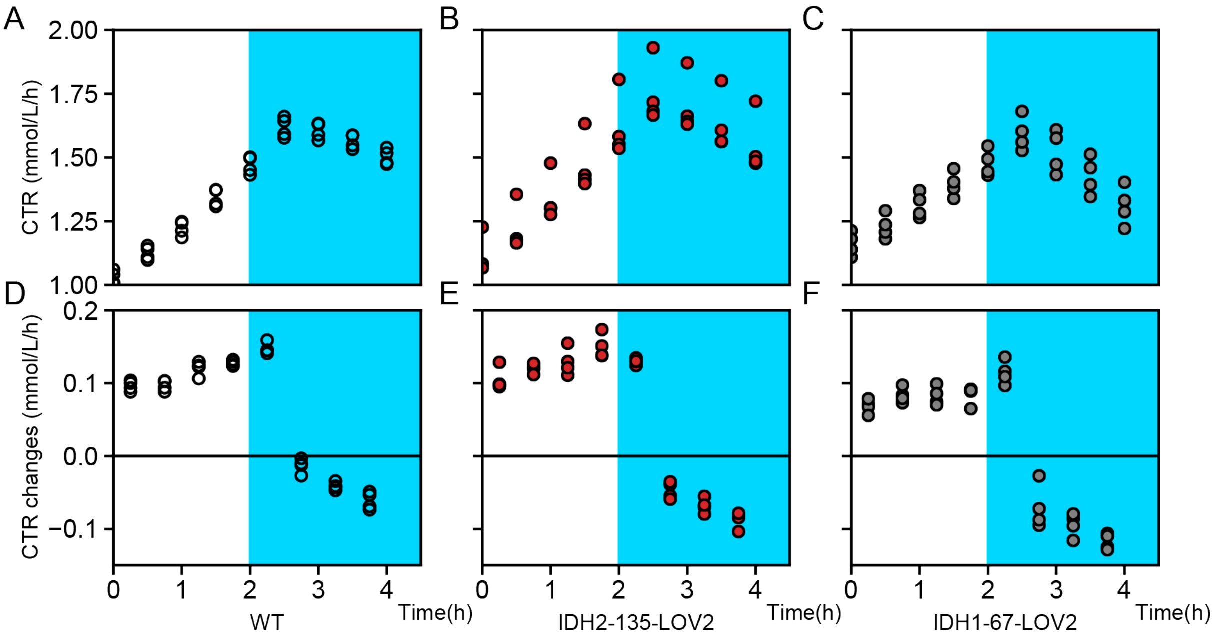


**Fig. S8: CTR measurement and response of light-switchable candidates. Related to Fig. 3.** (A-C) $CO_2$ transfer rate (CTR) measurements over time for WT (A), IDH2-135 (B) and IDH1-67 (C) strains. CTR were measured by the TOM device every 30 minutes. Blue shading indicates the phase of blue light illumination (470 nm, 750 $\mu W/cm^2$), starting from when the CTR measurement reached around 1.5 $mmol \cdot L^{-1} \cdot h^{-1}$. Four biological replicates were performed for each strain in YNB medium supplied with 1% acetate. After an initial slight CTR increase, likely due to a temperature increase caused by the LED light excitation, and LED temperature stabilization (Fig. S7D), CTR started decreasing as a result of illumination. (D-F) Momentary changes between two consecutive CTR measurements. Values from the second time points after turning on the blue light were used as the response to blue light illumination (Fig. 3C).

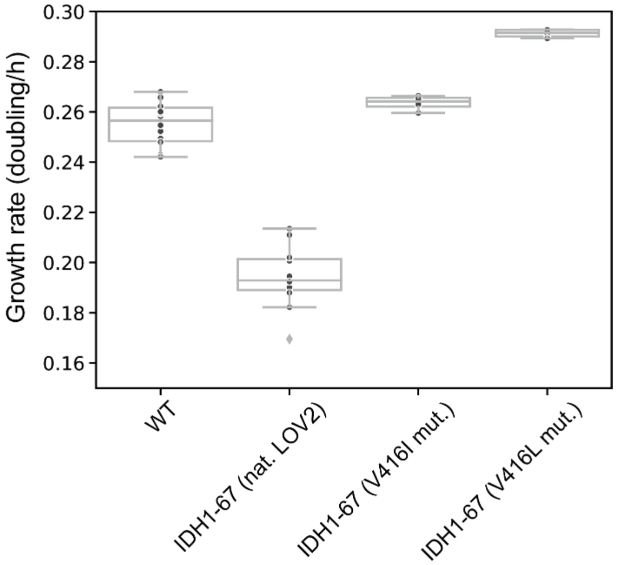


**Fig. S9. Growth rate measured from TOM assays.** The strains with slow-reverting LOV2 mutant showed a higher growth rate, estimated from the CTR measurements and shown as doubling per hour, than the strain with the natural LOV2 domain. Growth rates of WT, IDH1-67 (nat LOV2), IDH1-67 (V416I mut.) and IDH1-67 (V416L mut.) are estimated from 14, 11, 4 and 4 biological replicates, respectively.

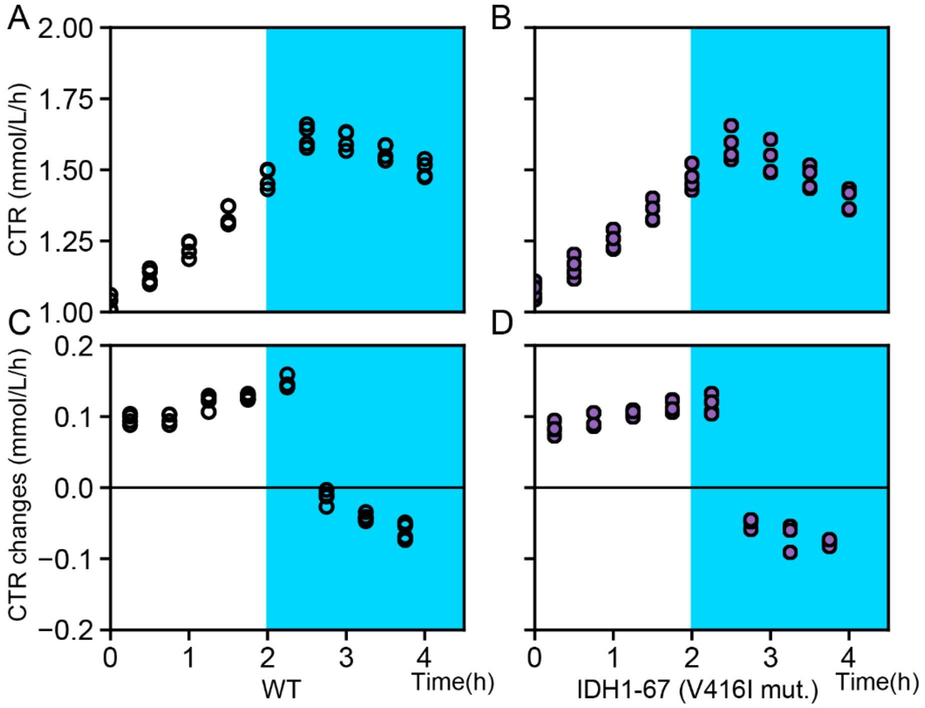


**Fig. S10: Related to Fig. 4.** (A-B) $CO_2$ transfer rate (CTR) measurements over time for WT (A, same as in Fig. S6A), IDH1-67 (V416I mut.) (B) strains. CTR were measured by the TOM device every 30 minutes. Blue shading indicates the phase of blue light illumination (470 nm, 750 $\mu W/cm^2$), starting from when the CTR measurement reached around 1.5 $mmol \cdot L^{-1} \cdot h^{-1}$. Four biological replicates were performed for each strain in YNB medium supplied with 1% acetate. After LED temperature stabilization (Fig. S5D), CTR started decreasing as a result of illumination. (C-D) Momentary changes between two consecutive CTR measurements for WT (C, same as in Fig. S6D) and IDH1-67 (V416I mut.) (D). Values from the second time points after turning on the blue light were used as the response to blue light illumination (Fig. 4D) strains.

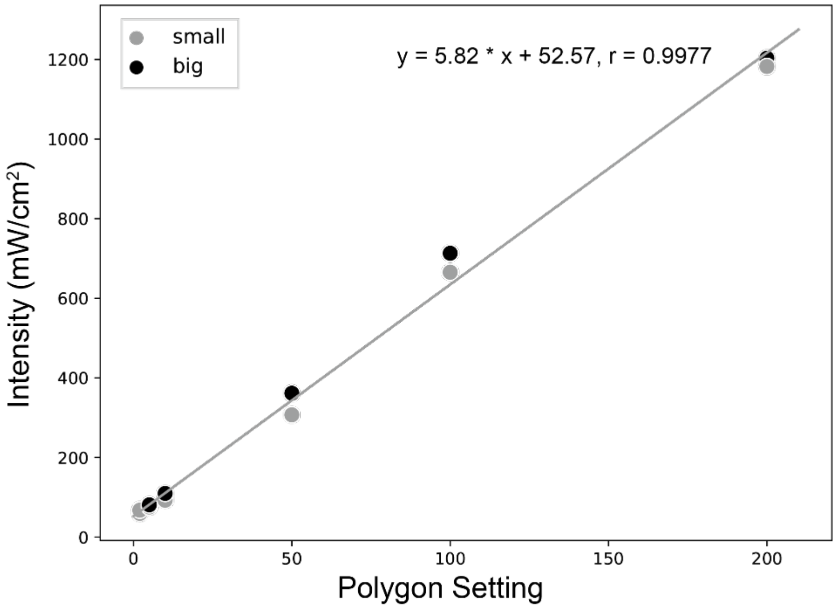


**Fig. S11: Intensity of Polygon lamp.** Intensity is measured by putting the light sensor S120VC on top of the objective to measured the power of light beam and divided by the area of the light beam, constrained by a pinhole in the light path. Results shown here are for configuration where no neutral density filter is applied and a 40x objective is used. During the experiment, a neutral density filter with OD=2 or OD=3 was applied to decrease the light intensity by 100- or 1000- folds (Fig. 5A, B and Fig. 12A, B, C), so that they fall within the range for optogenetics illumination. Results obtained with a large and a small light path pinhole are similar. A linear regression is fitted to estimate intensity at settings that are not measured.

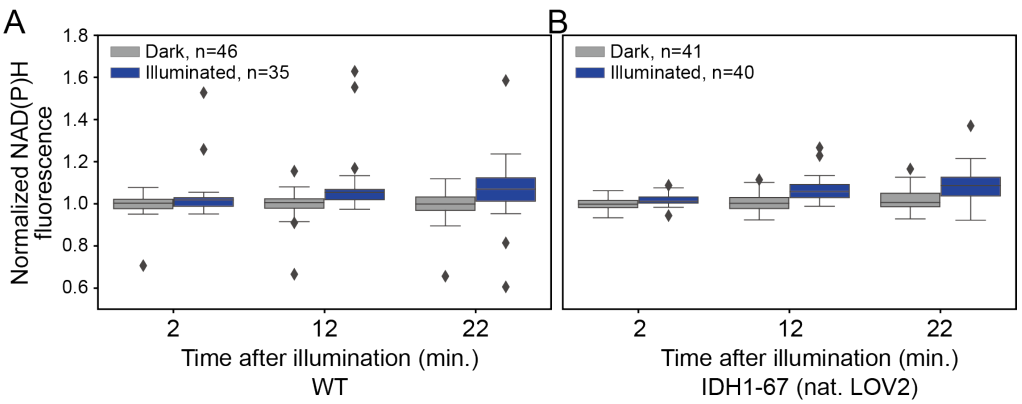


**Fig. S12. NAD(P)H response to blue light of lower light intensity.** NAD(P)H response of (A) Wildtype and (B) IDH1-67(natural LOV2) strains to blue light illumination. Blue light at an intensity of around 40 mW/cm2 was turned on two minutes before the first time point for the lit field of view. The NADH values of each cell are normalized to its value in the frame right before illumination.

**Supplementary Table** 1 **|** List of insertion sites and yeast strains

| **Variant ID*** | **Amino acid residue insertion** | **Insertion point (PDB: 3blx)** | **Priority (A=highest)** | **Remark from expert-eye inspection on the contact map and structure** | **solvent exposure residues (Å$^2$)** | **Summary of contacts (sites on 3blx)** |
|---|---|---|---|---|---|---|
| **IDH2-135-LOV2** | E-LOV2-G | 120-122 | A | Potential interactions via an antiparallel stretch into the active site. Not sure if LOV2 will fit here. | 124, 57, 30 | **AS**, contacting 135 and 248-259; **ABS**, contacting 138 -139; **AAR**, none |
| **IDH2-158-LOV2** | S-LOV2-G | 143 | B | In close proximity to active site. Possibly will work by affecting local structure. | 44 | AS, within 140-143, contacting 248-249; ABS, none, AAR, contacting 145-162 |
| **IDH2-160-LOV2** | I-LOV2-E | 145-147 | B | Same as above | 48, 37, 32 | **AS**, contacting 248-249. **ABS**, none, **AAR**, within145 to 162. |
| **IDH2** | - | 117 | C | Inspection suggests that introduction of LOV2 will not be feasible here. | 111 | same as for IDH2-135-LOV2 |
| **IDH1-67-LOV2** | Q-LOV2-T | 56-58 | B | Insertion straight into an allosteric control region. | 133, 38, 76 | **AS**, none; **ABS**, contacting 25-28 and 82-99; **AAR**, inside 53-60 |
| **IDH1-93-LOV2** | H-LOV2-T | 82 | B | Insertion straight into an allosteric control region. | 53 | **AS**, none; **ABS**, inside 82-99, contacting 25-28, 272-287, and IDH2-187-195; **AAR**, contacting 53-60 |
| **IDH1-185-LOV2** | N-LOV2-R | 174-175 | A | No problems were noticed when the structure was inspected. | 115,44 | **AS**, contacting 217-225; **ABS**, none; **AAR**, none |
| **IDH1-218-LOV2** | P-LOV2-D | **207**-208 | A/B | Contacts perhaps less convincing based on the map. Visual inspection indicates that an insertion could still give sufficient change in the local backbone to shift the global equilibrium between tense and relaxed state. | 81, 97 | **AS**, contacting 217-225; **ABS**, none; **AAR**, none |
| **IDH1-237-LOV2** | K-LOV2-P | 226 | A/B | Same as above | 109 | **AS,** next to 217-225, contacting 180-183, IDH2-248-259; **ABS,** contacting 241 and 245; **AAR**, none |

| | | | | | | |
|---|---|---|---|---|---|---|
| **IDH1-239-LOV2** | H-LOV2-Q | 228-229 | A/B | Same as above | 102,99 | same as above |
| **IDH1-275-LOV2** | R-LOV2-D | 264-265 | B | This position has a contact map that looks promising as an antiparallel stretch extends into an area involved in allosteric regulation. Unfortunately, the position is quite buried. | 138, 47 | **AS**, none, **ABS**, contacts with 82-99 and 272-287; **AAR**, none |
| **IDH1-288-LOV2** | G-LOV2-L | 277 | B | In close proximity to the AMP binding site. Therefore, insertion here could give too much disruption to the structure. | 45 | **AS**, none; **ABS**, within 272-287; **AAR**, none |
| **IDH1-291-LOV2** | I-LOV2-K | 280-281 | B | Same as above | 44, 158 | same as for IDH1-288-LOV2 |
| **IDH1-294-LOV2** | Q-LOV2-N | 283-284 | B | Same as above | 135, 49 | same as for IDH1-288-LOV2 |
| **IDH1** | - | 16-17 | C | It is unlikely that the inserted LOV2 domain would fit. | 33,89 | **AS**, none; **ABS**, contacting 79-99; **AAR**, contacting 53-60 |
| abbreviations: AS, active site; ABS, allosteric binding site for AMP and citrate; AAR, allosterically altered regions, where the protein changes conformation upon AMP and citrate binding but outside the binding site. | | | | | | |

*Numbers in the Variant IDs denote amino acid residue position on the full coding sequence (IDH1/IDH2, obtained from Saccharomyces Genome Database), before LOV2 domain insert.

**Supplementary Table 2 |** List of strains used in this study.

| Strain | Relevant genotype | Parental strain | Origin |
|---|---|---|---|
| IMX585 (referred to as WT in main text) | Can1Δ::Cas9-natNT2 | CEN.PK113-7D | Mans et al. [2] |
| IMX585 ΔIDH1 | Can1Δ::Cas9-natNT2 ΔIDH1 | IMX585 | This study |
| IMX585 ΔIDH2 | Can1Δ::Cas9-natNT2 ΔIDH2 | IMX585 | This study |
| IMX585 ΔIDH1ΔIDH2 | Can1Δ::Cas9-natNT2 ΔIDH1ΔIDH2 | IMX585 | This study |
| IDH1-67 | Can1Δ::Cas9-natNT2<br>IDH1 p.67Q_68TinsLOV2 | IMX585 | This study |
| IDH1-93 | Can1Δ::Cas9-natNT2<br>IDH1 p.93H_94TinsLOV2 | IMX585 | This study |
| IDH1-185<br>(with three non-synonymous mutations) | Can1Δ::Cas9-natNT2<br>IDH1 p.185N_186RinsLOV2, p.192V>L, p.193H>Y, p.194K>Q | IMX585 | This study |
| IDH1-185 | Can1Δ::Cas9-natNT2<br>IDH1 p.185N_186RinsLOV2 | IMX585 | This study |
| IDH1-218 | Can1Δ::Cas9-natNT2<br>IDH1 p.218P_219DinsLOV2 | IMX585 | This study |
| IDH1-237 | Can1Δ::Cas9-natNT2<br>IDH1 p.237K_238PinsLOV2 | IMX585 | This study |

| | | | |
|---|---|---|---|
| IDH1-239 | Can1Δ::Cas9-natNT2<br>IDH1 p.239H_240QinsLOV2 | IMX585 | This study |
| IDH1-275 | Can1Δ::Cas9-natNT2<br>IDH1 p.275R_276DinsLOV2 | IMX585 | This study |
| IDH1-288 | Can1Δ::Cas9-natNT2<br>IDH1 p.288G_289LinsLOV2 | IMX585 | This study |
| IDH1-291 | Can1Δ::Cas9-natNT2<br>IDH1 p.291I_292K | IMX585 | This study |
| IDH1-294 | Can1Δ::Cas9-natNT2<br>IDH1 p.294Q_295NinsLOV2 | IMX585 | This study |
| IDH2-135<br>(with three non-synonymous mutations) | Can1Δ::Cas9-natNT2<br>IDH2 p.135E_136GinsLOV2, p.140T>I, p.141Y>C, p.142E>A | IMX585 | This study |
| IDH2-135 | Can1Δ::Cas9-natNT2<br>IDH2 p.135E_136GinsLOV2 | IMX585 | This study |
| IDH2-158 | Can1Δ::Cas9-natNT2<br>IDH2 p.158S_159GinsLOV2 | IMX585 | This study |
| IDH2-160 | Can1Δ::Cas9-natNT2<br>IDH2 p.160I_161EinsLOV2 | IMX585 | This study |
| IDH1-67-lit | Can1Δ::Cas9-natNT2<br>IDH1 p.67Q_68TinsLOV2$_{I510E_I539E}$ | IMX585 | This study |
| IDH1-67-dark | Can1Δ::Cas9-natNT2 | IMX585 | This study |

| | | | |
|---|---|---|---|
| | IDH1 p.67Q_68TinsLOV2$_{C450M}$ | | |
| IDH2-135-lit | Can1Δ::Cas9-natNT2<br>IDH2<br>p.135E_136GinsLOV2$_{I510E_I539E}$ | IMX585 | This study |
| IDH2-135-dark | Can1Δ::Cas9-natNT2<br>IDH2 p.135E_136GinsLOV2$_{C450M}$ | IMX585 | This study |
| IDH1-67<br>(V416I mut.) | Can1Δ::Cas9-natNT2<br>IDH1 p.67Q_68TinsLOV2$_{V416I}$ | IMX585 | This study |
| IDH1-67<br>(V416L mut.) | Can1Δ::Cas9-natNT2<br>IDH1 p.67Q_68TinsLOV2 $_{V416L}$ | IMX585 | This study |
| IDH2-135<br>(V416I mut.) | Can1Δ::Cas9-natNT2<br>IDH2 p.135E_136GinsLOV2 $_{V416I}$ | IMX585 | This study |
| IDH2-135<br>(V416L mut.) | Can1Δ::Cas9-natNT2<br>IDH2 p.135E_136GinsLOV2 $_{V416L}$ | IMX585 | This study |

**Supplementary Table 3 | List of primers used in this study. sgRNA target sequence (20bp) indicated in red.**

| **Primer** | **Sequence** | **Functionality** |
|---|---|---|
| IDH1_targetRNA | TGCGCATGTTTCGGCGTTCGAAAC<br>TTCTCCGCAGTGAAAGATAAATG<br>ATCATGACAATCAAATCTATGTC<br>GTTTTAGAGCTAGAAATAGCAAG<br>TTAAAATAAG | IDH1 deletion |
| IDH1_repair oligo fw | GGTTCATTATTCTCCCTATCCTCA<br>TTCTTCTCCCTTTTCCTCCATAATT<br>GTAAGAGAAAATGAAAACAATTC<br>CCCTTTTTTTTGTTCTGCAACTTA<br>AGTGTGTTCAAATTCTTATCTATA | IDH1 deletion |
| IDH1_repair oligo rv | TATAGATAAGAATTTGAACACAC<br>TTAAGTTGCAGAACAAAAAAAAG<br>GGGAATTGTTTTCATTTTCTCTTA<br>CAATTATGGAGGAAAAGGGAGAA<br>GAATGAGGATAGGGAGAATAATG<br>AACC | IDH1 deletion |
| IDH2_targetRNA | TGCGCATGTTTCGGCGTTCGAAAC<br>TTCTCCGCAGTGAAAGATAAATG<br>ATCTCATTACTGAAGATAAAAGT<br>GTTTTAGAGCTAGAAATAGCAAG<br>TTAAAATAAG | IDH2 deletion |
| IDH2_repair oligo fw | ACCTTGGTAGATAATAAGATACA<br>GATCGGGAACGAACAACAATTAT<br>AATATTTTTAATAAAGTCCTATT<br>CTTTTCCCTCTCAGGTTTTTTCAC<br>GCCTTGAAAACAAATGACTATCC<br>GTT | IDH2 deletion |
| IDH2_repair oligo rv | AACGGATAGTCATTTGTTTTCAAG<br>GCGTGAAAAAACCTGAGAGGGAA<br>AAGAATAGGACTTTATTAAAAAA<br>TATTATAATTGTTGTTCGTTCCCG<br>ATCTGTATCTTATTATCTACCAAG<br>GT | IDH2 deletion |
| IDH1_targetRNA_#4 | TGCGCATGTTTCGGCGTTCGAAAC<br>TTCTCCGCAGTGAAAGATAAATG<br>ATCAAGCAAACAGATCATAAGGA<br>GTTTTAGAGCTAGAAATAGCAAG<br>TTAAAATAAG | IDH1-67 insertion |
| IDH1-67_repair | AGAACCATTTTTGAGGCTGAAAA | IDH1-67 insertion for the following |

| | | |
|---|---|---|
| oligo fw | TATCCCGATCGACTGGGAAACTATAAACATTAAGCAATTGGCTACTACACTTGAACG | strain:<br>IDH1-67<br>IDH1-67-lit<br>IDH1-67-dark<br>IDH1-67 (V416I mut.)<br>IDH1-67 (V416L mut.) |
| IDH1-67_repair oligo rv | ACCAATCTTATTTCTCTTTAGAGACTCAACAGCTTCATAGACGCCTTCTTTGTGGTCTGTAAGTTCTTTTGCCGCCTC | IDH1-67 insertion for the following strain:<br>IDH1-67<br>IDH1-67-lit<br>IDH1-67-dark<br>IDH1-67 (V416I mut.)<br>IDH1-67 (V416L mut.) |
| IDH1_targetRNA_#10 | TGCGCATGTTTCGGCGTTCGAAACTTCTCCGCAGTGAAAGATAAATGATCTCTCTAAAGAGAAATAAGATGTTTTAGAGCTAGAAATAGCAAGTTAAAATAAG | IDH1-93 insertion |
| IDH1-93_repair oligo fw | GTCTATGAAGCTGTTGAGTCTCTAAAGAGGAACAAAATTGGTCTTAAGGGGCTATGGCACTTGGCTACTACACTTGAACG | IDH1-93 insertion |
| IDH1-93_repair oligo rv | ATCTAGTTGTTTACGCAAAGCAACGTTTAGTGAACCGTGACCTGTTTGGTCAGCAGGAGTAAGTTCTTTTGCCGCCTC | IDH1-93 insertion |
| IDH1_targetRNA_#36 | TGCGCATGTTTCGGCGTTCGAAACTTCTCCGCAGTGAAAGATAAATGATCGTCTGTCACAGCTGTGCATAGTTTTAGAGCTAGAAATAGCAAGTTAAAATAAG | IDH1-185 insertion |
| IDH1-185_repair oligo fw | ACTAGACCTAAGACAGAAAGGATCGCCAGATTTGCCTTTGACTTCGCCAAGAAATACAACTTGGCTACTACACTTGAACG | IDH1-185 insertion |
| IDH1-185_repair oligo rv | GAACAGACCGTCACCTAACTTCATGATATTTGCCTGATACAGAGCTGTGACAGACTTTCTAAGTTCTTTTGCCGCCTC | IDH1-185 insertion (this primer introduces three non-synonymous mutations) |

| | | |
|---|---|---|
| IDH1-185-rep-rv-II | GAACAGACCGTCACCTAACTTCA TGATATTTGCCTTGTGTACTGCTG TGACAGACTTTCTAAGTTCTTTTG CCGCCTC | Repairing fragment construction of the following strain without non-synonymous mutations:<br><br>IDH1-185 |
| IDH1_targetRNA _#1 | TGCGCATGTTTCGGCGTTCGAAAC TTCTCCGCAGTGAAAGATAAATG ATCAGAAATATAATAACTGAAAT GTTTTAGAGCTAGAAATAGCAAG TTAAAATAAG | IDH1-218 insertion |
| IDH1-218_repair oligo fw | AAGTTAGGTGACGGTCTGTTCAG AAATATAATCACAGAGATTGGCC AAAAAGAATATCCTTTGGCTACT ACACTTGAACG | IDH1-218 insertion |
| IDH1-218_repair oligo rv | AGGTTTGGCCACCGCCTGCATGG AGGCATTGTCGACAATGATGGAC GATACGTCAATATCAAGTTCTTTT GCCGCCTC | IDH1-218 insertion |
| IDH1_targetRNA _#18 | TGCGCATGTTTCGGCGTTCGAAAC TTCTCCGCAGTGAAAGATAAATG ATCCTAGTTACCCCTTCAATGTAG TTTTAGAGCTAGAAATAGCAAGT TAAAATAAG | IDH1-237 insertion<br>IDH1-239 insertion |
| IDH1-237_repair oligo fw | CCTGATATTGACGTATCGTCCATC ATTGTCGACAATGCCTCCATGCA GGCGGTGGCCAAATTGGCTACTA CACTTGAACG | IDH1-237 insertion |
| IDH1-237_repair oligo rv | AATGTTGCCTAAGATGGTACCGT ACATAGATGGAGTAACTAGGACA TCAAATTGATGAGGAAGTTCTTTT GCCGCCTC | IDH1-237 insertion |
| IDH1-239_repair oligo fw | ATTGACGTATCGTCCATCATTGTC GACAATGCCTCCATGCAGGCGGT GGCCAAACCTCATTTGGCTACTAC ACTTGAACG | IDH1-239 insertion |
| IDH1-239_repair oligo rv | AGCGCCAATGTTGCCTAAGATGG TACCGTACATAGATGGAGTAACT AGGACATCAAATTGAAGTTCTTTT GCCGCCTC | IDH1-239 insertion |
| IDH1_targetRNA _#3 | TGCGCATGTTTCGGCGTTCGAAAC TTCTCCGCAGTGAAAGATAAATG | IDH1-275 insertion<br>IDH1-288 insertion |

| | | |
|---|---|---|
| | ATCAAACCAACATGTCTGGAACCGTTTTAGAGCTAGAAATAGCAAGTTAAAATAAG | IDH1-291 insertion<br>IDH1-294 insertion |
| IDH1-275_repair oligo fw | AACATTGGCGCTGCTTTGATCGGTGGTCCAGGATTGGTGGCAGGTGCCAACTTTGGCAGGTTGGCTACTACACTTGAACG | IDH1-275 insertion |
| IDH1-275_repair oligo rv | ATTTTGGCCTTTAATATCTAAACCAACATGCCTAGATCCTGGTTCGAAGACAGCATAGTCAAGTTCTTTTGCCGCCTC | IDH1-275 insertion |
| IDH1-288_repair oligo fw | GCAGGTGCCAACTTTGGCAGGGACTATGCTGTCTTCGAACCAGGATCTAGGCATGTTGGTTTGGCTACTACACTTGAACG | IDH1-288 insertion |
| IDH1-288_repair oligo rv | TAACGTGGAGGAAAGGATCATGGCAGTTGGGTTAGCCACATTTTGGCCTTTAATATCTAAAAGTTCTTTTGCCGCCTC | IDH1-288 insertion |
| IDH1-291_repair oligo fw | AACTTTGGCAGGGACTATGCTGTCTTCGAACCAGGATCTAGGCATGTTGGTTTAGATATTTTGGCTACTACACTTGAACG | IDH1-291 insertion |
| IDH1-291_repair oligo rv | GTTCAACATTAACGTGGAGGAAAGGATCATGGCAGTTGGGTTAGCCACATTTTGGCCTTTAAGTTCTTTTGCCGCCTC | IDH1-291 insertion |
| IDH1-294_repair oligo fw | AGGGACTATGCTGTCTTCGAACCAGGATCTAGGCATGTTGGTTTAGATATTAAAGGCCAATTGGCTACTACACTTGAACG | IDH1-294 insertion |
| IDH1-294_repair oligo rv | ACCCAAATGGTTCAACATTAACGTGGAGGAAAGGATCATGGCAGTTGGGTTAGCCACATTAAGTTCTTTTGCCGCCTC | IDH1-294 insertion |
| IDH2_targetRNA_#7 | TGCGCATGTTTCGGCGTTCGAAACTTCTCCGCAGTGAAAGATAAATGATCTAAATCAACGTTTTCGTAAGGTTTTAGAGCTAGAAATAGCAAGTTAAAATAAG | IDH2-135 insertion |
| IDH2-135_repair | TTGACATTGAGAAAAACATTTGG | IDH2-135 insertion for the |

| | | |
|---|---|---|
| oligo fw | GTTATTTGCCAACGTTCGTCCCGCAAAGTCTATTGAATTGGCTACTACACTTGAACG | following strains:<br>IDH2-135<br>DH2-135-lit<br>IDH2-135-dark<br>IDH2-135 (V416I mut.)<br>IDH2-135 (V416L mut.) |
| IDH2-135_repair oligo rv | ACCTTCGGTATTCTCTCTGATAAGAACTAAATCAACGTTTGCGCAAATGGTCTTAAAACCAAGTTCTTTTGCCGCCTC | IDH2-135 insertion for the following strains (this primer introduces three non-synonymous mutations):<br>IDH2-135<br>DH2-135-lit<br>IDH2-135-dark |
| IDH2-135-rep-rv-II | ACCTTCGGTATTCTCTCTGATAAGAACTAAATCAACGTTCTCATATGTGGTCTTAAAACCAAGTTCTTTTGCCGCCTC | IDH2-135 insertion for the following strains without non-synonymous mutations:<br>IDH2-135<br>DH2-135-lit<br>IDH2-135-dark<br>IDH2-135 (V416I mut.)<br>IDH2-135 (V416L mut.) |
| IDH2_targetRNA_#2 | TGCGCATGTTTCGGCGTTCGAAACTTCTCCGCAGTGAAAGATAAATGATCAATACCGAAGGTGAATATTCGTTTTAGAGCTAGAAATAGCAAGTTAAAATAAG | IDH2-158 insertion<br>IDH2-160 insertion |
| IDH2-158_repair oligo fw | ACCACTTACGAAAACGTTGATTTAGTTCTTATCAGAGAGAATACCGAAGGAGAGTACTCTTTGGCTACTACACTTGAACG | IDH2-158 insertion |
| IDH2-158_repair oligo rv | ATCTCTTGTGATCAGTTTAATAGATTGAACAACGCCAGGGCAAACTATGTGTTCGATACCAAGTTCTTTTGCCGCCTC | IDH2-158 insertion |
| IDH2-160_repair oligo fw | TACGAAAACGTTGATTTAGTTCTTATCAGAGAGAATACCGAAGGAGAGTACTCTGGTATCTTGGCTACTACACTTGAACG | IDH2-160 insertion |
| IDH2-160_repair | AGAGGCATCTCTTGTGATCAGTTT | IDH2-160 insertion |

| | | |
|---|---|---|
| oligo rv | AATAGATTGAACAACGCCAGGGCAAACTATGTGTTCAAGTTCTTTGCCGCCTC | |
| IDH1_dg fw | TGTGGGTGAGCACATAGGAC | IDH1 genome verification (deletion or Lov2 insert) |
| IDH1_dg rv | CAGCTGTCCAGAGTGAAGGG | IDH1 genome verification (deletion or Lov2 insert) |
| IDH1_seq fw | AGAGAACTTTAGCCACTGCC | IDH1 genome verification (Lov2 insert) |
| IDH1_seq fw2 | GAGAAAACACGGAGGGTGAG | IDH1 genome verification (Lov2 insert) |
| IDH1_seq rv | AGAAAAACCCCTGAGCAGAC | IDH1 genome verification (Lov2 insert) |
| IDH2_dg fw | AGGGGCCTTTTGCGATACTC | IDH2 genome verification (deletion or Lov2 insert) |
| IDH2_dg rv | GGTGAACATTGACTGCCAGC | IDH2 genome verification (deletion or Lov2 insert) |
| IDH2_seq fw | CATTCCTGACCCTGCCGTAC | IDH2 genome verification (Lov2 insert) |
| IDH2_seq rv | CAGATGGGTTGGTGACCACC | IDH2 genome verification (Lov2 insert) |
| LOV2-mid-fw | AGAAGGGAGATGTCCAGTAC | Sanger sequencing for all yeast strains with LOV2 |
| LOV2-mid-rv | CTTCACGGCTATATTCTGTC | Sanger sequencing for all yeast strains with LOV2 |
| LOV2-V416I/L-com-fw | GAATTATGCAGTGCTGCCATAAC | Gibson assembly to construct LOV2 V416I and V416L plasmids |
| LOV2-V416I/L-com-rv | GTTATGGCAGCACTGCATAATTC | Gibson assembly to construct LOV2 V416I and V416L plasmids |
| Lov2-V416I-fw | GAGAAGAACTTTATTATTACTGACCC | Gibson assembly to construct Plasmid V416I |
| Lov2-V416I-rv | GGGTCAGTAATAATAAAGTTCTTCTC | Gibson assembly to construct Plasmid V416I |
| LOV2-V416L-fw | GAGAAGAACTTTTTGATTACTGACCC | Gibson assembly to construct Plasmid V416L |
| Lov2-V416L-rv | GGGTCAGTAATCAAAAAGTTCTTCTC | Gibson assembly to construct Plasmid V416L |

| | | |
|---|---|---|
| mVenus-F | AACTCTCACAACGTCTACATC | Plasmid sequencing of all template plasmids. |

**Supplementary Table 4 | List of plasmids used in study**

| Name | Functions | Origin | Addgene |
|---|---|---|---|
| pROS13-IDH1 #1 | sgRNA for IDH1-218 insertion | This study | |
| pROS13-IDH1 #3 | sgRNA for IDH1-275, IDH1-288, IDH1-291, IDH1-294 insertion | This study | |
| pROS13-IDH1 #4 | sgRNA for IDH1-67 insertion | This study | 166101 |
| pROS13-IDH1 #10 | sgRNA for IDH1-93 insertion | This study | |
| pROS13-IDH1 #18 | sgRNA for IDH1-237, IDH1-239 insertion | This study | |
| pROS13-IDH1 #36 | sgRNA for IDH1-185 insertion | This study | |
| pROS13-IDH2 #2 | sgRNA for IDH2-158, IDH2-160 insertion | This study | |
| pROS13-IDH2 #7 | sgRNA for IDH2-135 insertion | This study | 166102 |
| Plasmid V416I | Template plasmid for LOV2 V416I mutant | This study | |
| Plasmid V416L | Template plasmid for LOV2 V416L mutant | This study | |
| pSwi4-mVenus-LOV2 | Template plasmid for LOV2 domain | This study | |
